# Colibactin-induced genotoxicity and colorectal cancer exacerbation critically depends on adhesin-mediated epithelial binding

**DOI:** 10.1101/2023.08.16.553526

**Authors:** Maude Jans, Magdalena Kolata, Gillian Blancke, Maarten Ciers, Anders B. Dohlman, Takato Kusakabe, Mozes Sze, Alexandra Thiran, Geert Berx, Sabine Tejpar, Geert van Loo, Iliyan D. Iliev, Han Remaut, Lars Vereecke

**Affiliations:** VIB Center for Inflammation Research, B-9052 Ghent, Belgium; Department of Biomedical Molecular Biology, Ghent University, B-9052 Ghent, Belgium; Department of Internal Medicine and Pediatrics, Ghent University, Ghent, Belgium; Ghent Gut Inflammation Group, Ghent University, Ghent, Belgium; Cancer Research Institute Ghent (CRIG), 9000 Ghent, Belgium; Structural Biology Brussels, Vrije Universiteit Brussel, Brussels, Belgium; Structural & Molecular Microbiology, VIB-VUB Centre for Structural biology, Brussels, Belgium; Department of Medical Oncology, Dana-Farber Cancer Institute, Boston, MA, USA; Joan and Sanford I. Weill Department of Medicine, Weill Cornell Medicine, Cornell University, New York, NY 10021, USA; The Jill Roberts Institute for Research in Inflammatory Bowel Disease, Weill Cornell Medicine, NY 10065, USA; Department of Oncology, University Hospital Leuven, KU Leuven, Leuven, Belgium

**Keywords:** Colorectal cancer, colibactin, pks, Escherichia coli, microbiome, genotoxin, epithelial adhesion, DNA damage

## Abstract

Various bacteria are suggested to contribute to colorectal cancer (CRC) development, including *pks+ E. coli* which produce the genotoxin colibactin that induces characteristic mutational signatures in host epithelial cells. It remains unclear how the highly unstable colibactin molecule is able to access host epithelial cells and its DNA to cause harm. Using the microbiota-dependent ZEB2-transgenic mouse model of invasive CRC, we found that *pks+ E. coli* drives CRC exacerbation and tissue invasion in a colibactin-dependent manner. Using isogenic mutant strains, we further demonstrate that CRC exacerbation critically depends on expression of the *E. coli* type-1 pilus adhesin FimH and the F9-pilus adhesin FmlH. Blocking bacterial adhesion using a pharmacological FimH inhibitor attenuates colibactin-mediated genotoxicity and CRC exacerbation. Together, we show that the oncogenic potential of *pks+ E. coli* critically depends on bacterial adhesion to host epithelial cells and is critically mediated by specific bacterial adhesins. Adhesin-mediated epithelial binding subsequently allows production of the genotoxin colibactin in close proximity to host epithelial cells, which promotes DNA damage and drives CRC development. These findings present promising therapeutic avenues for the development of anti-adhesive therapies aiming at mitigating colibactin-induced DNA damage and inhibiting the initiation and progression of CRC, particularly in individuals at risk for developing CRC.

## Introduction

Approximately 2 million patients worldwide are diagnosed with colorectal cancer (CRC) annually. With approximately half of these patients succumbing to the disease, CRC ranks as the third most prevalent and the third deadliest form of cancer^1^. CRC incidence is associated to Western lifestyle and is increasing, particularly in young individuals (age younger than 50)^2,3^. An increasing body of research provides strong evidence pointing to the indispensable involvement of the gut microbiota in the initiation and progression of CRC, operating through direct or indirect mechanisms that include the disruption of the intestinal epithelial barrier, detrimental inflammation, immune dysregulation, and genotoxicity^4^. The concept of bacterial-host interactions playing a significant role in CRC development gains additional support through the discovery of various CRC-promoting or ‘onco’bacteria, including *Fusobacterium nucleatum*, enterotoxigenic *Bacteroides fragilis* and pathogenic *Escherichia coli* strains^5–8^. While the majority of *E. coli* strains are harmless commensals, the species includes several pathotypes that cause acute diarrheal disease, as well as strains linked to chronic infections and conditions such as inflammatory bowel disease (IBD) and even CRC^9^. Among these strains are ‘adherent-invasive *E. coli*’ (AIEC) strains, which possess the ability to attach to and invade host cells^10,11^. Several *E. coli* strains are known to induce DNA damage by producing a set of enzymes encoded in the ‘polyketide synthase’ pathogenicity island (*pks*), which synthesize the genotoxin colibactin. These *pks+ E. coli* strains have been detected in approximately 20% of healthy individuals, around 40% of patients with IBD, and approximately 60% of CRC patients, and show an association with the inflamed and neoplastic mucosa, where they are thought to contribute to a crucial positive feedback mechanism in CRC onset and progression^12–14^. Colibactin possesses hydrophilic warheads capable of binding adenine residues, crosslinking DNA and inducing double strand breaks^15–17^.

*In vitro* infection studies have demonstrated that *pks+ E. coli* can cause cell cycle arrest and double-strand DNA breaks, while *ex vivo* infection of colonic loops in healthy mice has shown a significant increase in DNA damage to colonic epithelial cells^18,19^. Furthermore, human organoids injected with *pks+ E. coli* acquired a distinct mutational signature that has also been detected in a subset of CRC patients^20^. Using a synthetic colibactin molecule^21^, designed to mimic the bioactivity of natural colibactin, Dougherty and colleagues observed an increase in the total number of mutations, along with an increased occurrence of mutational signatures associated with mismatch repair deficiency (MMRd), suggesting that colibactin may exacerbate MMRd-associated mutations^22^. It is worth noting that the genotoxic effects of colibactin require direct contact between bacteria and host cells, as they are attenuated when bacteria are separated from mammalian cells using a cell-impermeable membrane^18^. However, fundamental knowledge on how colibactin can access host nuclei to inflict DNA damage is lacking.

In this study, we investigated the carcinogenic potential of *pks+ E. coli* and its role in promoting CRC development *in vivo*, using our recently developed microbiota-dependent mouse model of invasive CRC, based on transgenic expression of Zeb2 (Zeb2^IEC-Tg/+^). Transgenic expression of Zeb2, a transcription factor regulating epithelial-to-mesenchymal transition (EMT), in intestinal epithelial cells leads to spontaneous invasive colorectal cancer, which is completely prevented in germ-free conditions^23^. Therefore, Zeb2^IEC-Tg/+^ mice provide a unique preclinical model to investigate the underlying mechanisms through which *pks+ E. coli* may promote CRC development *in vivo*.

## Results

### *E. coli* genes are enriched in human CRC biopsies and *E. coli 11G5* infection exacerbates tumor development in Zeb2^IEC-Tg/+^ mice

Pathogenic E. coli strains have been associated with the inflamed and neoplastic mucosa of colorectal cancer (CRC) patients^12–14^. We analyzed the presence of *E. coli* specific sequences in The Cancer Genome Atlas (TCGA) sequencing data and found that their prevalence is increased in colon tumors compared to matched normal adjacent tissue (NAT) (Fig. 1A). Multiple *E. coli* genes were significantly more prevalent in CRC tumor tissue compared to normal adjacent tissue (NAT), including *yhbQ*, a DNA damage response nuclease (Fig. 1B), indicating that *E.coli* is preferentially accumulating in tumor tissue.

**FIGURE 1.**
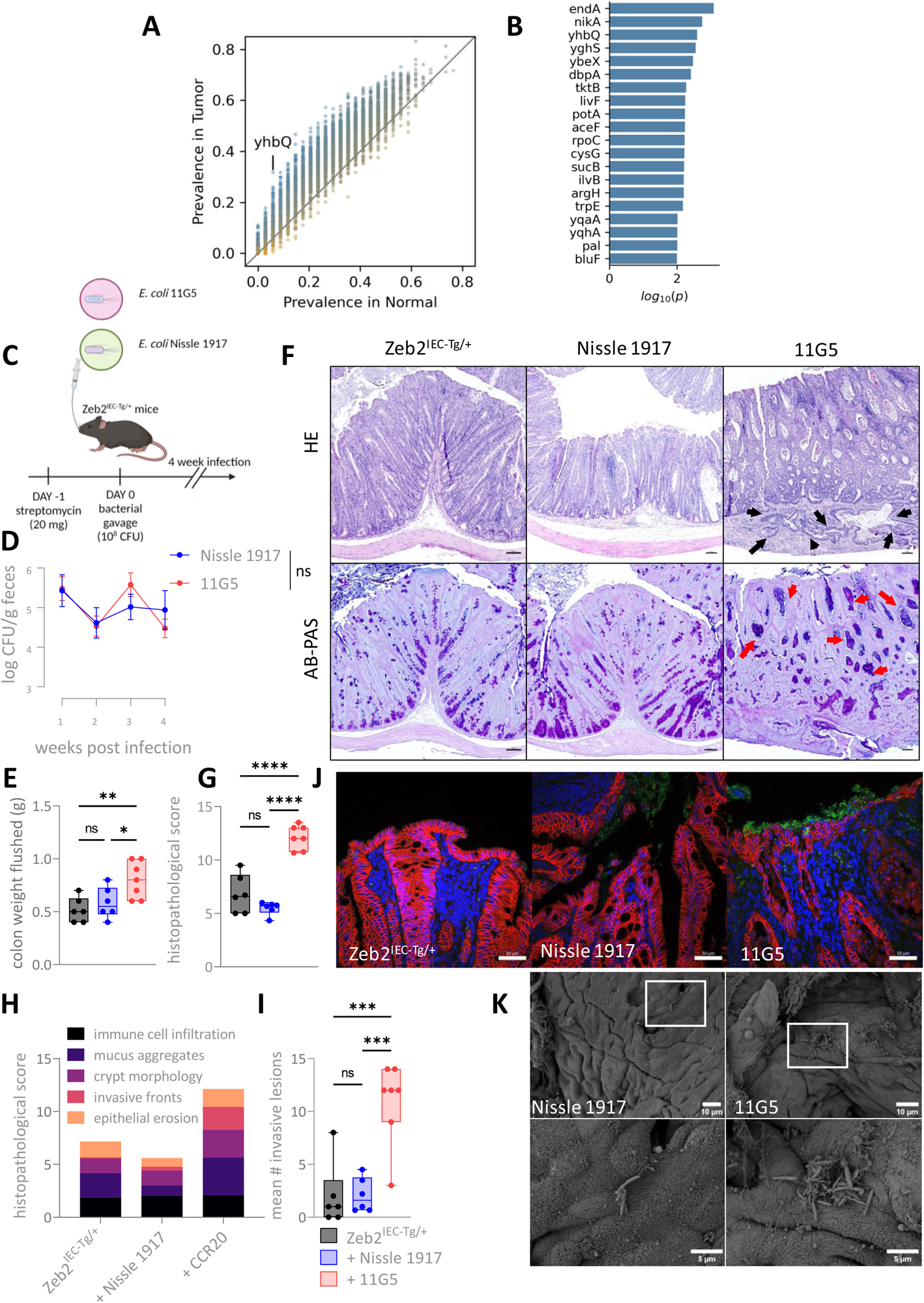
*E. coli 11G5* infection exacerbates tumor development in Zeb2^IEC-Tg/+^ mice and induces a DNA damage response. **A.** Prevalence of *E. coli* genes in TCGA sequencing data from The Cancer Microbiome Atlas (TCMA) from CRC patients versus normal adjacent tissue (NAT). **B.** *E. coli* genes significantly enriched in human CRC tumor tissue, determined by Fisher’s exact test. **C.** Experimental timeline of infections. Mice received streptomycin pretreatment followed by oral gavage of 10^8^ CFU *E. coli Nissle 1917* or *E. coli 11G5*. Mice were sacrificed after a 4-week infection. **D.** Fecal colonisation of *E. coli Nissle 1917* and *E. coli 11G5* monitored during the course of the infection. Data are represented as mean ± SEM. **E.** Flushed colon weights of Zeb2^IEC-Tg/+^ mice after *Nissle 1917* (n = 6) and *11G5* (n = 7) infection or PBS control (n = 6). Data are represented as mean ± SEM. (* p < 0.05, ** p < 0.01; 1-way ANOVA with multiple comparisons). **F.** Representative images of hematoxylin eosin (H&E) (upper panel)-, Alcian blue/periodic acid–Schiff (AB-Pas) (lower panel) stained colon sections of Zeb2^IEC-Tg/+^ mice after *E. coli Nissle 1917* and *E. coli 11G5* infection or non-infected controls (PBS). Black arrows indicate epithelial invasive lesions into the submucosa, red arrows indicate mucus aggregates. Scale bar: 100 μm. **G.** Histopathological score of tumor sections following *11G5* and *Nissle 1917* infection. Data are represented as mean ± SEM. (**** p < 0.0001; one-way ANOVA with multiple comparisons). **H.** Histopathological score indicating parameters affected by infection. **I.** Mean number of invasive lesions into the submucosa. Data are represented as mean ± SEM. (*** p < 0.001; one-way ANOVA with multiple comparisons). **J.** Representative images of *E. coli* (green) and E-cadherin (red) staining on tumor sections of uninfected, *Nissle 1917*- or *11G5*-infected Zeb2^IEC-Tg/+^ mice following a 4-week infection. Scale bar 50μm. **K.** scanning electron microscopy (SEM) images of colons of Zeb2^IEC-Tg/+^ mice following Nissle 1917 or 11G5 infection. Scale bar overview: 10 μm; scale bar zoom (lower panel): 5 μm.

To elucidate whether *pks+ E. coli* affects tumor development *in vivo*, we made use of the Zeb2^IEC-Tg/+^ mouse model of microbiota-dependent CRC, which we developed previously. Zeb2^IEC-Tg/+^ mice express the EMT-regulator Zeb2 specifically in intestinal epithelial cells^23^, and are characterized by a destabilized epithelial and mucus barrier, which promotes bacterial-epithelial interactions, inflammation and spontaneous microbiota dependent invasive CRC-development^23^. Therefore, Zeb2^IEC-Tg/+^ mice are the ideal experimental model to study the complex microbe-host interplay during CRC development, particularly during colonization with suspected ‘oncobacteria’ such as *pks+ E. coli*. We infected Zeb2^IEC-Tg/+^ mice with *E. coli 11G5*, a human *pks+* CRC isolate (previously also referred to as *E. coli* CCR20, personal communication – Richard Bonnet)^24^. 6-week old specific-pathogen-free (SPF) Zeb2^IEC-Tg/+^ mice were orally gavaged with streptomycin to overcome colonization resistance. After 24h, mice were orally challenged with a single dose of 10^8^ CFU of *E. coli 11G5 (*hereafter referred to as *‘11G5’)*, with the commensal strain *E. coli Nissle 1917 (*hereafter *‘Nissle 1917’),* or with PBS (control) (Fig. 1C). Quantification of fecal CFUs throughout the experiment revealed similar levels of *Nissle 1917* and *11G5*, indicating that downstream effects are not caused by differences in colonisation efficacy of either strain (Fig. 1D). After 4-weeks infection, mice were sacrificed for histopathological analysis. *11G5*-colonized Zeb2^IEC-Tg/+^ mice showed increased colon weight, suggesting a higher tumor burden (Fig. 1E). To evaluate tumor progression at the histological level, we developed a histopathological scoring system to quantify the typical histopathological features of Zeb2^IEC-Tg/+^ tumors, taking into account immune cell infiltration, mucus accumulation, epithelial erosion, invasive lesions and crypt morphology. Tumors of Zeb2^IEC-Tg/+^ mice infected with *11G5* showed exacerbated tumor development with a significant increase in the histopathological score, which is characterized by increased mucus aggregates (red arrows) and tissue invasion (black arrows) (Fig. 1F-H). Quantification of tissue invasion revealed an increase in the mean number of epithelial invasive lesions into the submucosa in tumors of *11G5*-infected mice compared to control- and *Nissle 1917*-infected mice (Fig. 1I).

To verify whether *E. coli 11G5* promotes tumor development either directly by affecting epithelial cells, or indirectly through immune-mediated effects, we evaluated multiple inflammation parameters. *11G5* infection did not affect fecal lipocalin2 levels, a commonly used parameter of intestinal inflammation (Supplementary Fig. 1A). Increased levels of IL-17A, IL-1β and CXCL1 could be detected in the serum of *11G5*-infected mice compared to *Nissle 1917*-infected mice, while no differences in concentration of proinflammatory cytokines other than IL-17A could be observed in *11G5*-infected mice compared to PBS-treated control Zeb2^IEC-Tg/+^ mice (Supplementary Fig. 1B). Additionally, transcriptome analysis based on bulk RNA-sequencing of sorted CD45+ colonic lamina propria immune cells after either *E. coli Nissle 1917* or *E. coli 11G5* infection in Zeb2^IEC-Tg/+^ mice indicated a similar expression profile of the immune compartment, with only 10 differentially expressed genes (8 downregulated, 2 upregulated) (Supplementary Fig. 1C,D). Taken together, these data indicate that *E. coli 11G5* does not strongly change immune activation in Zeb2^IEC-Tg/+^ mice, which suggests that its tumor promoting functions are unlikely to be attributed to indirect immune-related effects.

To examine possible direct effects of *E. coli 11G5* on the colonic epithelium, we visualized bacterial epithelial interactions by performing immunofluorescence staining for *E. coli* (green) and E-cadherin (epithelial cells, red) on tumor samples of *E. coli 11G5-* or *Nissle 1917*-infected versus uninfected Zeb2^IEC-Tg/+^ mice. In uninfected and *Nissle 1917*-infected mice, *E. coli* could only be detected in the gut lumen. In contrast, we found that *11G5* strongly associated with the colonic surface epithelium and also invaded the epithelial layer and underlying lamina propria of Zeb2^IEC-Tg/+^ mice (Fig. 1J, Supplementary Fig. 1E). Using scanning electron microscopy (SEM) on colonic segments of Zeb2^IEC-Tg/+^ mice after infection with either *11G5* or *Nissle 1917*, we observed more and larger *E. coli*-shaped bacterial colonies associated to the intestinal epithelium in *11G5*-infected mice (Fig. 1K). Additionally, we observed enriched epithelial surface-associated and crypt-associated *11G5* compared to *Nissle 1917* both in proximal and distal colonic segments, despite similar fecal colonisation (Supplementary Fig. 1F).

Since *E. coli 11G5* adheres to the intestinal epithelium and thrives in this microenvironment, we evaluated the transcriptional expression profile of sorted epithelial cells from Zeb2^IEC-Tg/+^ mice following either *E. coli Nissle 1917* or *11G5* infection. Principal component analysis (PCA) revealed distinct epithelial expression profiles/transcriptomes (Fig. 2A), with a greater number of genes being downregulated in *11G5* versus *Nissle 1917* control group (Fig. 2B). Genes upregulated in epithelial cells after *11G5* infection were associated with various cellular processes, including epithelial-to-mesenchymal transition (EMT), cellular invasion, tumor-promoting pathways including Notch1, p53 and E2F1 signaling, metabolism and cell cycle (Fig. 2C, Supplementary Fig. 1G). Interestingly, *11G5* infection also activated pathways related to DNA repair and DNA damage bypass (Fig. 2D). In contrast, *11G5* infection in WT background did not induce morphological changes, intestinal inflammation or changes in the transcriptional expression profile of both epithelial cells and CD45+ immune cells (Supplementary Fig. 1H-K). Taken together, these data indicate that *pks+ E. coli 11G5* closely associates with the colonic epithelium and exacerbates CRC development in Zeb2^IEC-Tg/+^ mice. *E. coli 11G5* does not induce profound changes in immune activation but is associated with increased DNA damage-response mechanisms in colonic epithelial cells.

**FIGURE 2.**
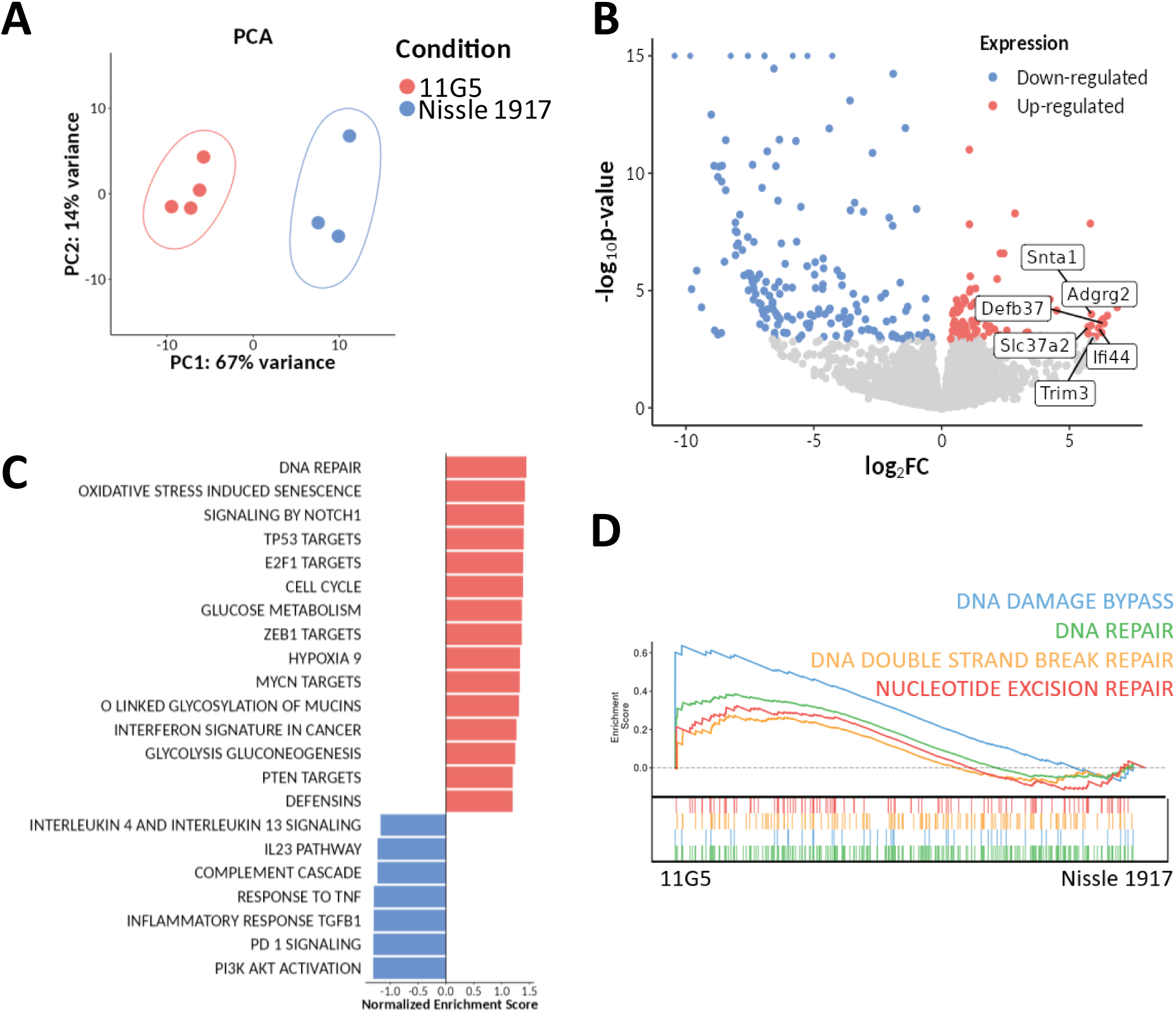
*E. coli 11G5* induces transcriptional changes in intestinal epithelium of Zeb2IEC-Tg/+ mice and instigates a DNA damage response. **A.** Principal component analysis (PCA) showing distinct overall gene expression profiles in epithelial cells sorted from the colon of Zeb2^IEC-Tg/+^ mice following 4-week *Nissle 1917* (n = 3) or *11G5* (n = 4) infection. **B.** Volcano plot of differentially expressed genes (DEG) in colonic epithelial cells of *11G5* infected Zeb2^IEC-Tg/+^ mice compared to *Nissle 1917*. **C.** Gene set enrichment analysis (GSEA) results of pathways related to cancer development. FC, fold-change. **D.** Enrichment score for reactome pathways related to DNA damage, DNA repair and DNA damage bypass.

### Tumor-promoting activity of *11G5* is colibactin-dependent

Since DNA damage response and repair pathways are upregulated in epithelial cells of *E. coli 11G5*-infected Zeb2^IEC-Tg/+^ mice compared to *E. coli Nissle 1917*-infected mice, we next explored the role of colibactin on the enhanced tumorigenesis observed in Zeb2^IEC-Tg/+^ mice colonized with *E. coli 11G5*. The genotoxin colibactin is produced by a set of enzymes encoded in the *pks* pathogenicity islands, and genetic deletion of the essential thioesterase *ΔclbQ* prevents colibactin production^25^. We infected Zeb2^IEC-Tg/+^ mice with either wild-type (WT) *11G5* or with the colibactin-deficient isogenic mutant strain *11G5ΔclbQ* that no longer produces active colibactin^24,26^. Since *Nissle 1917* also harbors the *pks* pathogenicity islands, we also included the non-genotoxic *Nissle 1917ΔclbQ* mutant as control strain. Fecal CFU counts of these strains was shown to be similar to the wild-type strain, and *E. coli 11G5ΔclbQ* was shown to adhere to the colonic epithelium to a similar extent as the wild-type strain (Supplementary Fig. 2A,B). Remarkably, CRC exacerbation could no longer be observed in Zeb2^IEC-Tg/+^ mice colonized with *E. coli 11G5ΔclbQ*, as shown by significantly reduced colon weight, histology, histopathological scoring and level of tumor invasion in *11G5ΔclbQ*-infected mice, compared to mice infected with WT *11G5* (Fig. 3A-D). Although *E. coli Nissle 1917* contains *pks* pathogenicity islands and is believed to produce colibactin, we observed no difference in tumor development in Zeb2^IEC-Tg/+^ mice infected with either WT or *clbQ*-deficient *E. coli Nissle 1917* strains (Fig. 3A-D). Given the genotoxic potential of colibactin, we examined phosphorylation of histone H2Ax (γH2Ax), a biomarker of double stranded DNA breaks and DNA repair^27^. For this, we stained formalin-fixed paraffin-embedded (FFPE) tumor samples for γH2Ax and Epcam, to visualize and quantify DNA damage in colonic epithelial cells (Fig. 3E,F). *E. coli Nissle 1917* infection did not induce DNA damage responses in the colonic epithelium of infected Zeb2^IEC-Tg/+^ mice, as γH2Ax staining was not increased compared to uninfected Zeb2^IEC-Tg/+^ mice or mice infected with *E. coli Nissle 1917ΔclbQ* (Fig. 3E,F). In contrast, however, WT *11G5* infection, but not infection with the *11G5ΔclbQ* isogenic mutant strain, significantly increased the number of γH2Ax+ epithelial cells per crypt compared to uninfected and *E. coli Nissle 1917*-infected Zeb2^IEC-Tg/+^ mice (Fig. 3E,F). In agreement, *in vitro* infection of the colorectal cell line HT-29 with *11G5* showed strong DNA damage responses, which could not be observed upon infection with the *11G5ΔclbQ* strain, as shown by western blot analysis for γH2Ax. *Nissle 1917* and *Nissle 1917ΔclbQ* infection did not induce DNA damage in HT-29 cells (Fig. 3G). Taken together, these data demonstrate that *E. coli 11G5* promotes CRC development in Zeb2^IEC-Tg/+^ mice through colibactin-mediated genotoxic effects. Remarkably, despite containing the *pks* pathogenicity island, *E. coli Nissle 1917* does not promote measurable DNA damage or CRC development *in vivo*, and has only mild genotoxic effects *in vitro*.

**FIGURE 3.**
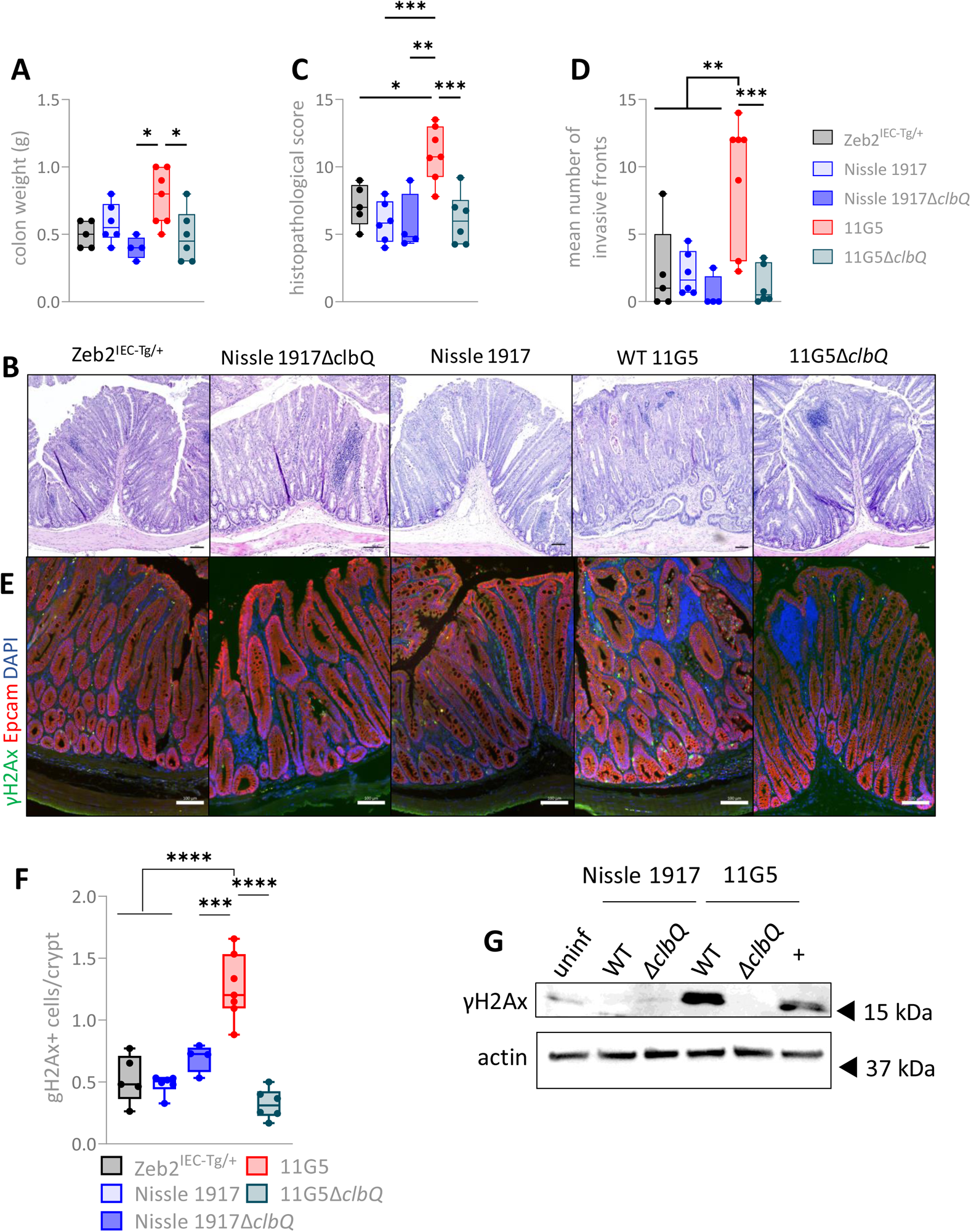
Tumor-promoting activity of *11G5* is colibactin-dependent. **A.** Flushed colon weights of Zeb2^IEC-Tg/+^ mice, uninfected (n = 5) or infected with *Nissle 1917* (n = 6), *Nissle 1917ΔclbQ* (n = 4), *11G5* (n = 7) or *11G5ΔclbQ* (n = 6). Data are represented as mean ± SEM. (* p < 0.05; one-way ANOVA with multiple comparisons). **B.** Representative images of H&E-stained colon sections of Zeb2^IEC-Tg/+^ mice following a 4 week infection with *Nissle 1917*, *11G5* or isogenic mutant strains (*ΔclbQ)*. Scale bar: 100 μm. **C & D.** Histopathological score and mean number of invasive lesions after a 4-week infection respectively. Data are represented as mean ± SEM. (* p < 0.05, ** p <0.01, *** p<0.001; one-way ANOVA with multiple comparisons). **E.** Representative images of γH2Ax (green) and Epcam (red) immunostaining on tumor sections. Scalebar overview, 100 μm; scalebar zoom, 50 μm. **F.** Quantification of the number of γH2Ax-positive epithelial cells in tumor sections of Zeb2^IEC-Tg/+^ mice after infection. Data are represented as mean ± SEM. (*** p<0.001, **** p<0.0001; one-way ANOVA with multiple comparisons). **G.** HT-29 cells were infected with indicated *E. coli* strains for 6h. Immunoblot analysis for γH2Ax 24h after infection. Positive control (+): HT-29 cells treated with 1 μM staurosporine. Actin was used as loading control.

### Carcinogenic effects of pks+ *E. coli* depend on adhesin-mediated epithelial binding

Since *E. coli 11G5,* but not *pks+ E. coli Nissle 1917,* can drive colibactin-mediated CRC exacerbation, we reasoned that additional virulence factors must contribute to the carcinogenic properties of *11G5*. Colibactin is a labile molecule whose genotoxic activity has been found to be contact-dependent^18,28^. Host cell adherence may therefore be a strong determinant for the genotoxic capacity of *pks*+ *E. coli*. To evaluate the importance of epithelial binding for *E. coli 11G5*’s oncogenic functions, we investigated two prominent bacterial adhesins expressed by *11G5*: the type 1 (*fim*) and F9 (*fml*) pilus adhesins FimH and FmlH, which respectively bind terminal D-mannose in oligomannose N-glycans, and the Thomsen-Friedenreich (TF) antigen^29,30^. The type 1 pilus is well known for its ability to drive the formation of extra- and intracellular bacterial communities (or biofilms) of uro- and enteropathogenic *E. coli* (including AIEC) in bladder and gut^31–33^. The F9 pilus is of interest since mucosal glysolyation alterations occur in precancerous conditions (UC, CD) and colon cancer, thus exposing the TF antigen^34,35^. We generated *11G5 fimH* and *fmlH* isogenic mutant strains (*11G5ΔfimH* and *11G5ΔfmlH*) and performed infection experiments in Zeb2^IEC-Tg/+^ mice. Six-week old Zeb2^IEC-Tg/+^ mice were infected with either WT *11G5* or with the adhesin mutants for a period of 4 weeks, after which we evaluated their capacity to bind the colonic epithelium *in vivo*. In contrast to WT *11G5*, which closely adheres to the colonic epithelium, both adhesin-mutant strains were mostly present in the intestinal lumen and were not in close contact with the colonic mucosa (Fig. 4A), although no differences in bacterial colonisation of the intestinal tract were observed between *11G5*-, *11G5ΔfimH-* and *11G5ΔfmlH*-infected mice (Supplementary Fig. 3A-C). Subsequent histological analysis revealed a significant reduction in tumor severity in mice infected with the adhesin mutants compared to mice infected with the WT *11G5* strain (Fig. 4B,C). In addition, significantly less invasive lesions into the submucosa were observed in tumors of mice infected with the adhesin mutants compared to WT *11G5* (Fig. 4D). To further investigate the epithelial cell interaction of WT *11G5* versus the isogenic *fimH* and *fmlH* mutant strains, we performed scanning electron microscope (SEM) analysis of HT-29 cells after infection with the commensal *E. coli Nissle 1917*, WT *11G5,* the colibactin-deficient *11G5ΔclbQ* or the adhesin mutant *11G5ΔfimH* and *11G5ΔfmlH* strain. Only WT *11G5* strongly associated to the cell surface of HT-29 cells and showed cell invasion (Fig. 4E). While Nissle 1917 showed minimal epithelial adhesion, both FimH and FmlH-adhesin knockout strains failed to bind epithelial HT-29 cells *in vitro*, and only showed adherence to the culture plate plastic (Fig. 4E). In contrast, the colibactin deficient *11G5ΔclbQ* strain was shown to be adhesion-competent and was shown to still associate to the cell membrane (Supplementary Fig. 3D). We also quantified epithelial adhesion *in vitro* using flow-cytometry, by infecting HT-29 and HCT116 cells with CFSE-stained *E. coli*. In both cell lines, adhesin mutants *11G5ΔfimH* and *11G5ΔfmlH* could no longer adhere to epithelial cells, in contrast to WT *11G5* (Fig. 4F, Supplementary Fig. 3E-G).

**FIGURE 4.**
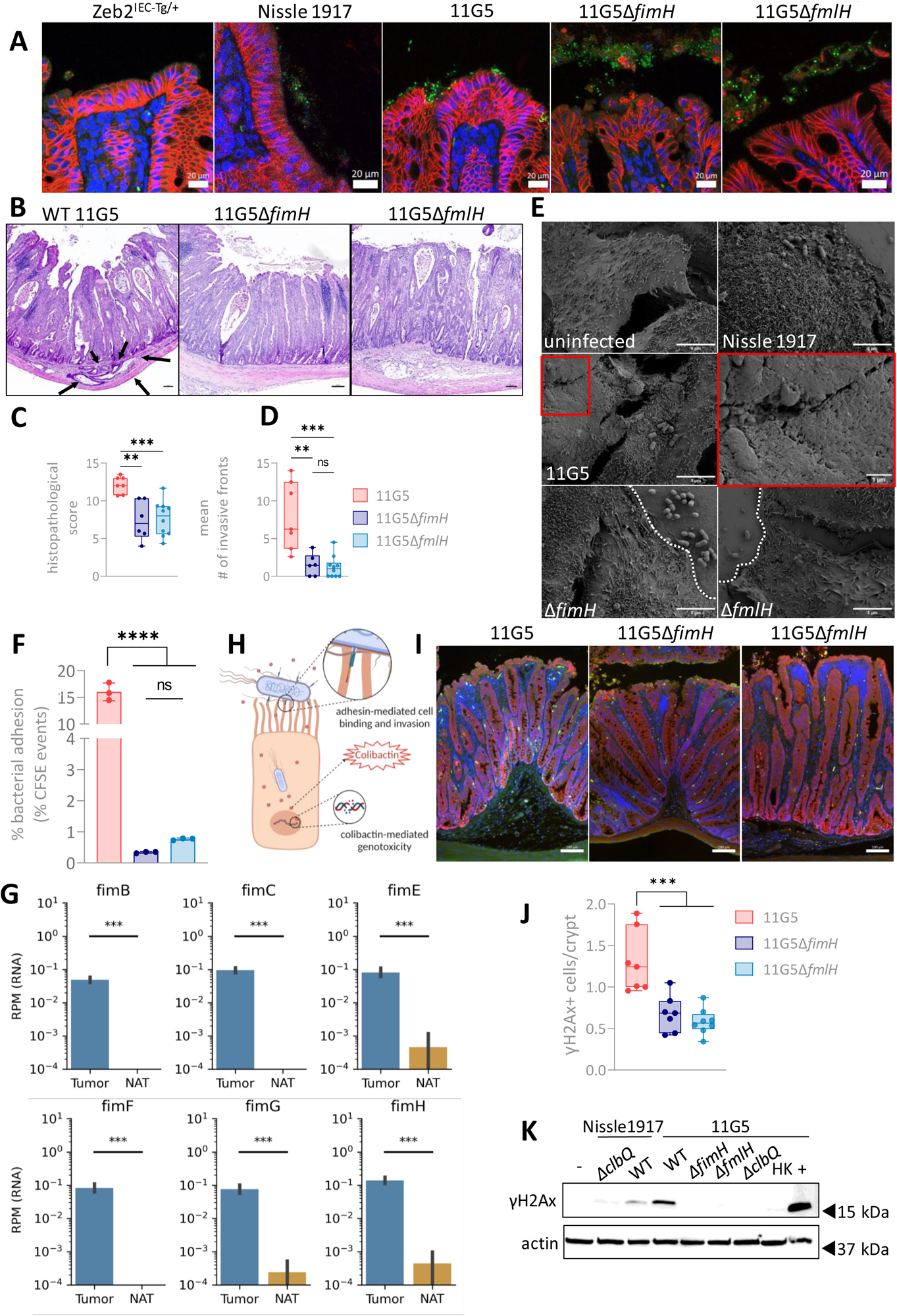
Effect of *11G5* on tumor development is dependent on FimH- and FmlH-mediated adhesion to the colonic epithelium. **A.** Representative images of *E. coli* (green) and E-cadherin (red) immunostaining of tumor sections of uninfected, *Nissle 1917*, *11G5* or isogenic adhesin mutant (*11G5ΔFimH*, *11G5ΔFmlH*) infected Zeb2^IEC-Tg/+^ mice following a 4-week infection. Scale bar: 20μm. **B.** Representative images of H&E-stained colon sections of Zeb2^IEC-Tg/+^ mice following a 4 week infection with WT *11G5* (n = 7) or isogenic mutants *11G5ΔfimH* (n = 6) and *11G5ΔfmlH* (n = 10). Black arrows indicate epithelial invasive lesions. Scale bar: 100 μm. **C.** Histopathological score of Zeb2^IEC-Tg/+^ mice infected with WT *11G5*, *11G5ΔfimH* or *11G5ΔfmlH*. Data are represented as mean ± SEM. (** p <0.01, *** p<0.001; one-way ANOVA with multiple comparisons). **D.** Mean number of invasive lesions into the submucosa. Data are represented as mean ± SEM. (** p <0.01, *** p<0.001; one-way ANOVA with multiple comparisons). **E.** Representative scanning electron microscopy (SEM) images of HT-29 cells infected with respective *E. coli* strains. Border of epithelial cells with culture plate plastic is denoted with white dashed line. Infection of 3h. Scale bar: 5 μm; scale bar zoom: 1 μm. **F.** Flow cytometric analysis of bacterial adhesion using HCT116 cells infected with CFSE-stained WT *11G5*, *11G5ΔfimH* or *11G5ΔfmlH*. Cells were infected for 3h and washed 3 times with PBS to remove loosely adherent bacteria. Data are represented as mean ± SD. Data is representative for two independent experiments. (** p <0.01, *** p < 0.001; one-way ANOVA with multiple comparisons). **G.** Abundance of RNA transcripts aligning to genes from the *E. coli* Fim cluster, including FimH, in human CRC tumors and adjacent NAT tissue in the TCGA cohort (*** p < 0.001). **H.** Schematic representation of the proposed hypothesis in which adhesin-mediated epithelial binding is required for colibactin-mediated DNA damage. **I.** Representative images of γH2Ax (green) and Epcam (red) immunostaining of tumor sections of Zeb2^IEC-Tg/+^ mice infected with WT *11G5*, *11G5ΔfimH* or *11G5ΔfmlH* 28 days after infection. Scale bar: 100 μm. **J.** Quantification of the number of γH2Ax-positive epithelial cells in tumor sections of Zeb2^IEC-Tg/+^ mice after infection. Data are represented as mean ± SEM (*** p<0.001; one-way ANOVA with multiple comparisons). **K.** Western blot analysis for γH2Ax in HT-29 cells infected with *E. coli Nissle 1917*, *11G5* or isogenic mutants. Cells were infected for 6h and medium was replaced with gentamicin-containing medium for 18h. HK: heat-killed, positive control (+): cells treated with 1 μM staurosporine.

Through the analysis of microbial RNA in TCGA sequencing data, we have previously identified the presence of transcriptionally active *Candida* species in CRC tumors^36^. Employing a similar approach, we now also detected higher expression of multiple genes from the *fim* cluster including *fimH* in human CRC tumors in contrast to non-tumorigenic tissue (Fig. 4G). Collectively, these findings demonstrate that the exacerbation of CRC mediated by *11G5* is heavily relying on bacterial-epithelial adhesion, showing an important contribution of both the type-1 pilus adhesin FimH and the F9 pilus adhesin FmlH. Furthermore, our data analysis of human samples indicate that adhesion-related genes like *fimH* are enriched in human CRC tumors.

### Colibactin-mediated genotoxicity critically depends on bacterial-epithelial adhesion

Since *E. coli 11G5* requires expression of both FimH and FmlH adhesins but also expression of the genotoxin colibactin to drive CRC progression, we hypothesized that the unstable colibactin molecule only acts at short-range distance, and can only exert genotoxic effects when it is produced in close proximity to host cell DNA (Fig. 4H). To investigate whether exposure to *E. coli 11G5* versus isogenic mutants inflicts DNA damage, we examined the presence of γH2Ax in the colonic epithelium of Zeb2^IEC-Tg/+^ mice after 4 weeks of infection. In line with our previous results, we found increased γH2Ax+ epithelial cells per crypt in tumors of Zeb2^IEC-Tg/+^ mice infected with *11G5* (Fig. 4I,J). However, an infection with adhesin mutants *ΔfimH* and *ΔfmlH* abrogated the genotoxic effect of *11G5* and no longer induced DNA damage in colonic crypts of Zeb2^IEC-Tg/+^ mice (Fig. 4I,J). To further test the involvement of bacterial adhesion on colibactin-mediated DNA damage, HT-29 cells were infected for 6h with *E. coli* strains (multiplicity of infection, MOI of 100), and γH2Ax was analysed 24h after infection by western blotting. We found increased γH2Ax levels in HT-29 cells infected with WT *11G5*, but not after infection with the colibactin-deficient *ΔclbQ* mutant strain (Fig. 4K). Similarly, HT-29 cells infected with adhesin mutants *11G5ΔfimH* and *11G5ΔfmlH* did not show activation of the DNA damage response (Fig. 4K). HT-29 cells and HCT116 cells infected with WT *E. coli Nissle 1917* showed moderate γH2Ax levels, which were abrogated in *Nissle 1917ΔclbQ* (Fig. 4K, Supplementary Fig. 3H). These findings were confirmed with immunofluorescence analysis, as only HT-29 cells infected with WT *11G5* showed nuclear γH2Ax foci, but not cells infected with the *Nissle 1917* strain or with the mutant *11G5ΔfimH*, *11G5ΔfmlH* or *ΔclbQ* strain (Supplementary Fig. 3I). Together, these data demonstrate that colibactin-mediated genotoxicity and CRC exacerbation in Zeb2^IEC-Tg/+^ mice critically depends on bacterial adhesin-mediated epithelial binding.

### Anti-adhesive therapy prevents tumor-promoting activity of *pks+ E. coli 11G5*

Our findings that CRC critically depends on bacterial-epithelial adhesion open therapeutic perspectives, as anti-adhesin-based anti-bacterials could be effective strategies to prevent colibactin-mediated DNA damage in humans. Sibofimloc is a potent FimH adhesin blocker which was recently found to be safe and effective in patients with active Crohn’s disease^37^. Therefore, we investigated the potential of sibofimloc to inhibit *pks+ E. coli 11G5* adhesion and colibactin-mediated DNA damage. Flow cytometric analysis showed that sibofimloc could significantly reduce adhesion of WT *11G5* to HCT116 cells in a dose-dependent manner (Fig. 5A). Furthermore, treatment with sibofimloc was shown to suppress DNA damage of *11G5*-infected HCT116 cells in a dose-dependent manner, as shown by γH2Ax Western blotting (Fig. 5B).

**FIGURE 5.**
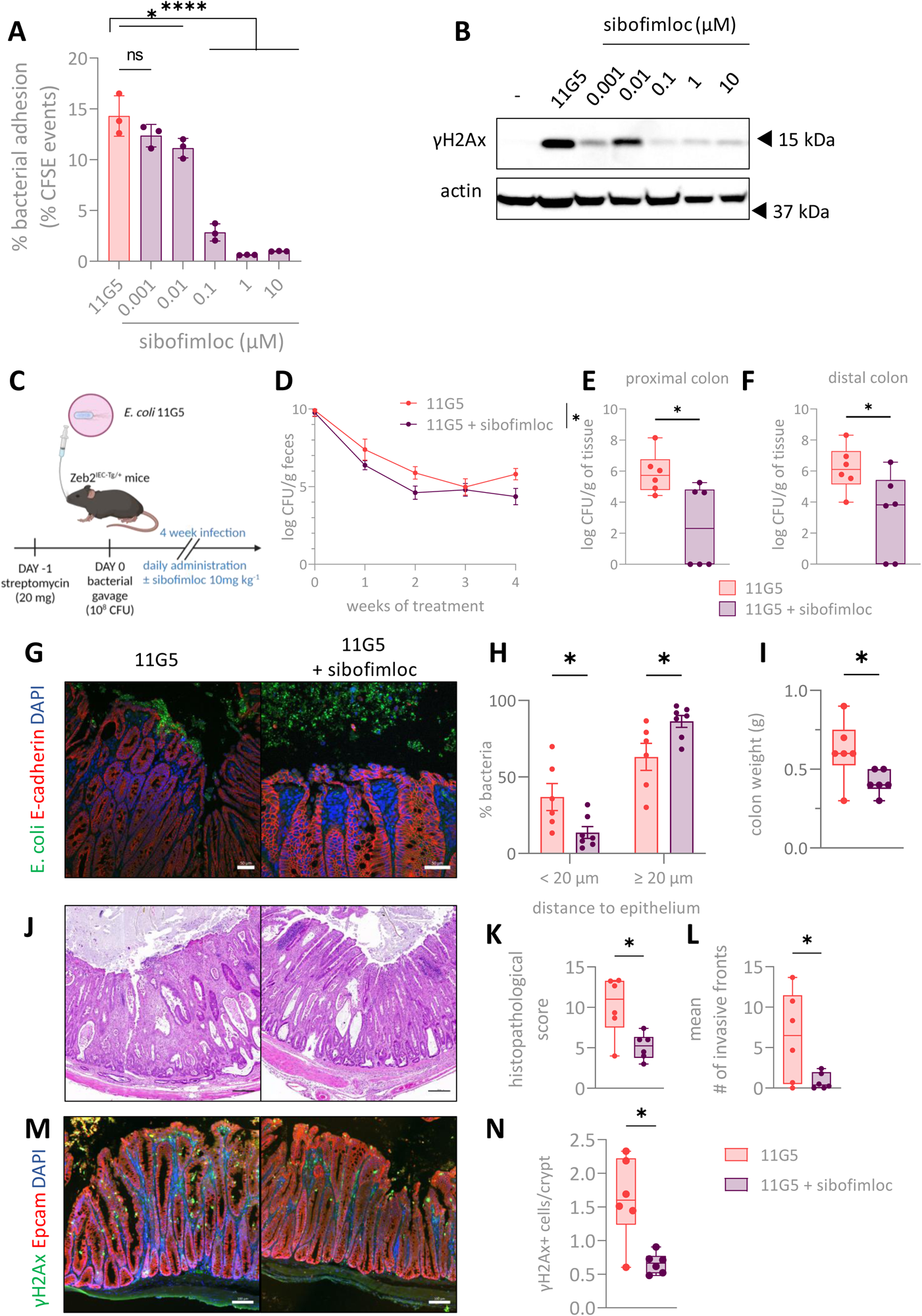
Anti-adhesive therapy abrogates colibactin-dependent tumor-promoting activity of 11G5. **A.** Flow cytometric analysis bacterial adhesion using HCT116 cells infected with CFSE-stained WT *11G5* with indicated sibofimloc concentrations. Data are represented as mean ± SD and is representative of two independent experiments. (* p < 0.05, **** p<0.0001; 1-way ANOVA with multiple comparisons). **B.** Western blot analysis of γH2Ax on HT-29 cells infected with *11G5* in presence of sibofimloc. Actin was used as loading control. Data is representative for three independent experiments **C.** Experimental set-up of therapeutic treatment with sibofimloc (or vehicle DMSO) following *11G5* infection. **D.** Fecal colonisation of 11G5 monitored over time. **E.** & **F.** CFU in proximal and distal colon homogenates upon sibofimloc (n = 6) or vehicle (n = 6) daily treatment respectively. (*p <0.05; two-sided unpaired t-test). **G.** Representative images of *E. coli* (green) and E-cadherin (red) immunostaining of tumor sections of. Scale bar: 50μm. **H.** Percentage of bacteria closely associated to the colonic epithelium (minimal average distance to epithelium < 20 μm) and not associated with the epithelium (minimal average distance to epithelium ≥ 20 μm) after 11G5 infection and treatment with sibofimloc or treatment. Data are represented as mean ± SEM. (*p < 0.05, two-way ANOVA). **I.** Flushed colon weights of 11G5-infected following sibofimloc treatment. Data are represented as mean ± SEM. (* p < 0.05, two-sided unpaired t-test). **J.** Representative images of H&E stained colon sections of Zeb2^IEC-Tg/+^ mice following a 4 week infection with *11G5* with sibofimloc treatment or vehicle-treated control. Scale bar: 200 μm. **K.** Histopathological score of Zeb2^IEC-Tg/+^ mice infected with WT *11G5*, followed by daily vehicle or sibofimloc treatment. Data are represented as mean ± SEM. (* p < 0.05; two-sided unpaired t-test). **L.** Mean number of invasive lesions into the submucosa. Data are represented as mean ± SEM. (* p < 0.05; two-sided unpaired t-test). **M.** Representative images of γH2Ax (green) and Epcam (red). Scale bar: 100 μm. **N.** Quantification of the number of γH2Ax-positive epithelial cells in tumor sections of Zeb2^IEC-Tg/+^ mice after infection. Data are represented as mean ± SEM (* p < 0.05; two-sided unpaired t-test with Welch’s correction).

Next, we studied the effects of blocking *11G5* adhesion with sibofimloc *in vivo*. Zeb2^IEC-Tg/+^ mice were infected with *11G5* followed 24h later by daily treatment with sibofimloc (10mg/kg body weight) or vehicle (DMSO), over the course of a four week infection period (Fig 5C). *11G5-* and sibofimloc-treated Zeb2^IEC-Tg/+^ mice showed significantly reduced *11G5* levels in feces and colon compared to those treated with a vehicle control, indicating that sibofimloc could effectively prevent epithelial binding *in vivo* (Fig 5D-F). Reduced epithelial binding was confirmed by anti-*E. coli* staining on colonic sections (Fig 5G-H). Sibofimloc was shown to displace *11G5* from the epithelial surface, suggesting strong affinity competition for FimH binding and neutralization (Fig 5G-H). Importantly, *in vivo* anti-adhesive therapeutic intervention using sibofimloc significantly reduced tumor burden, shown by decreased colon weight, histopathological score and tissue invasion in Zeb2^IEC-Tg/+^ mice (Fig 5I-L). In agreement, sibofimloc treatment significantly reduced epithelial DNA damage induced by *E. coli 11G5* (Fig 5M-N). Together, these data clearly demonstrate that pks+ *E. coli* rely on adhesin-mediated epithelial binding in order to allow colibactin-induced genotoxicity and CRC exacerbation.

Finally, we acquired freshly isolated intra-tumoral *E. coli* strains through the gentamicin protection assay (GPA) from lesional and healthy adjacent tissue biopsy samples from CRC patients. We isolated a novel *pks+ E. coli* strain (*MK54*) from a tumor biopsy, and investigated the effect of the *MK54 pks+ E. coli* CRC isolate on tumor development *in vivo* by infecting Zeb2^IEC-Tg/+^ mice with either *E. coli Nissle 1917* or *pks+ E. coli MK54*. After a 4-week infection, we again observed a significant increase in tumor burden, DNA damage in epithelial cells and bacterial adhesion to the colonic epithelium (Supplementary Fig 4A-H). We also evaluated the strain’s ability to adhere to epithelial cells and found that the *MK54 pks+ E. coli* CRC isolate could strongly adhere to HCT116 cells *in vitro*. *MK54* cell adherence was inhibited by sibofimloc treatment in a dose-dependent manner (Supplementary Figure 4I). Furthermore, we observed an increase in γH2Ax levels after 1 hour of infection (MOI 10) and up to 24h with the *pks+ E. coli* CRC isolate, which was dependent on epithelial binding, as sibofimloc treatment could reduce the DNA damage response in a dose-dependent manner (Supplementary Figure 4J).

In conclusion, our data provide new insights into how oncogenic *pks E. coli* strains contribute to CRC development, showing that adhesion-mediated epithelial association is crucial for colibactin-mediated DNA damage and CRC exacerbation. Importantly, our data demonstrate that specific adhesin-targeting strategies are effective to block CRC-promoting mechanisms.

## Discussion

Several bacteria have been identified as tumor-promoting bacteria or ‘oncobacteria’, such as *Fusobacterium nucleatum*, *enterotoxigenic Bacteroides fragilis* and *pks+ E. coli* strains^5–8,20^. *pks+ E. coli* have a tropism for the inflamed and neoplastic mucosa, as their abundance is increased in inflammatory bowel disease (IBD) and even more in colorectal cancer (CRC) patients^12–14^. Their increased presence in tumor biopsies compared to normal adjacent tissue also suggests that *pks+ E. coli* may drive critical feedback mechanisms for tumor development, possibly mediated through the synthesis of the genotoxin colibactin. Several studies have demonstrated that *pks+ E. coli* can promote colorectal cancer development in mice, as shown in the Apc^Min/+^ and in the AOM/DSS-induced tumor model, as well as in a chronic intestinal inflammation model ^11,24,38–41^. However, a thorough understanding of how colibactin mediates host damage or how colibactin translocates from bacteria to host cells is lacking^19,20^. Direct bacteria-host contact seems to be required for pathogenesis, since membrane separation of bacteria and mammalian cells prevents genotoxicity^18,42^. In this study, we investigated the relationship between epithelial adhesion of *pks+ E. coli* and tumor development. We demonstrate that expression of the chaperone-usher pili (CUP) adhesins FimH and FmlH is essential for colibactin-mediated genotoxicity. Remarkably, however, deletion of either FimH or FmlH offers significant protection from *pks+ E. coli*-induced pathology, without functional redundancy. The type-1 pilus adhesin FimH binds α-D-mannosylated proteins, while the F9-pilus adhesin FmlH binds Gal(β1-3)GalNac epitopes on O-glycoproteins, commonly termed the ‘Thomsen-Friedenreich’ (TF) antigen^29,34,35^. It remains unclear whether a broad range or rather specific epithelial glycoproteins are recognized by FimH and FmlH in the murine colon, or whether they act in synergy to ensure stable binding to host cells.

Various *E. coli* strains contain the *pks* pathogenicity island, including ‘beneficial’ strains which are used as probiotics, including *E. coli Nissle* 1917^43–46^. The idea that the probiotics currently in use possess the genetic capability to produce genotoxins is indeed concerning. However, our studies, using both the wild-type *pks+ E. coli Nissle* 1917 strain and an isogenic mutant *Nissle 1917* strain lacking clbQ, showed that neither strain promoted CRC development in Zeb2^IEC-Tg/+^ mice, indicating that, in contrast to more pathogenic strains like *11G5*, *E. coli Nissle* 1917 is much less oncogenic. Our *in vitro* genotoxicity assays demonstrate only mild γH2Ax staining upon *E. coli Nissle* 1917 infection, which was abrogated by *clbQ* loss. These observations indicate that, despite the presence of the *pks* pathogenicity island, *E. coli Nissle* 1917 has mild genotoxic potential, but may not be considered harmless. This difference in pathogenicity may be explained by either reduced cell-adhesive capacities of the *E. coli Nissle* 1917 strain, since adhesion to epithelial cells is considered a prerequisite for pathogenic bacteria. Indeed, i*n vitro* studies have demonstrated that IBD *E. coli* isolates adhere more frequently to intestinal tissue compared to *E. coli* isolates from healthy subjects^47–49^. An alternative explanation could be that *Nissle 1917* has reduced colibactin production potential compared to more pathogenic *pks+ E. coli* strains. However, deeming the utilization of *Nissle 1917* as a probiotic safe remains uncertain, given the potential for strain variation and evolutionary changes that might elevate its adhesive and toxigenic properties.

Our studies demonstrated a tumor-promoting effect of *E. coli 11G5*, a human *pks+* CRC isolate, on tumor development in Zeb2^IEC-Tg/+^ mice compared to uninfected or *E. coli Nissle 1917*-infected controls (both *pks+* and *pks-*). We hereby showed that tumor exacerbation by *pks+ E. coli 11G5* is mediated predominantly by *11G5*’s direct genotoxic effects on the host intestinal epithelial cells, rather than through indirect immune-mediated effects. In line with this, we demonstrated that the inflammatory status upon *E. coli 11G5* infection was not drastically affected.

Zeb2^IEC-Tg/+^ mice are characterized by an instable intestinal barrier and a defective colonic mucus layer, facilitating direct bacterial interactions^23^, which promote *pks+ E. coli* mediated toxicity. In contrast, in WT mice with intact mucosal barrier, *11G5* is unable to attach to the epithelium and induce tumorigenesis. A previous study demonstrated that even a thin mucus layer, as seen in the jejunum, is sufficient to attenuate the genotoxic effects of colibactin, suggesting that direct contact with the host epithelium is required for pathogenesis^42^. Destabilization of the colonic mucus layer therefore represents a significant risk for individuals colonized with *pks+ E. coli*. IBD patients typically display goblet cell depletion and mucus defects, and Muc2 levels are significantly lower in UC patients with active disease^50,51^. Also, a fiber-free Western-diet is known to promote degradation of the intestinal mucus layer, as several bacterial taxa shift to degradation of mucus-associated glycoproteins as a nutrient source in absence of fiber^52^. Therefore, intermittent or chronic fiber deprivation can promote mucus barrier defects and epithelial binding of *pks+ E. coli*, promoting CRC.

Importantly, we could show that pharmacological inhibition of FimH, using the small molecule FimH-blocker sibofimloc (Enterome), that has been approved for the treatment of Crohn’s disease, can prevent *pks+ E. coli*-mediated genotoxicity and CRC exacerbation. These finding unveil intriguing therapeutic possibilities for mitigating colibactin-induced DNA damage by targeting the inhibition of adhesin-mediated cell binding. Anti-adhesive therapies can be used to deplete or suppress selective bacterial species without affecting the resident microbiota and host physiology^31^.

Finally, our research extends beyond the specific *pks+ E. coli* strain *11G5*, as anti-adhesive therapy exhibited a comparable reduction in genotoxic potential on another CRC isolated *pks+ E. coli* strain associated with CRC. This suggests that the observed effects are not limited to a single pathogenic strain, but that anti-adhesive therapy could be an effective strategy against *pks+ E. coli* in general. Various strategies to target *pks+ E.coli* or colibactin synthesis have been reported. A recent study successfully targeted colibactin biosynthesis using a small molecule inhibitor of ClbP, the peptidase responsible for processing precolibactin to the mature genotoxin colibactin. This ClbP inhibitor was shown to effectively suppress colibactin production, even within a complex gut microbiome, without affecting other bacteria through antibiotic activity^53^. Hence, modulation of gut microbial functions through administration of small compounds emerges as a promising avenue for therapeutic intervention^53^. Multiple studies have also shown promising potential for targeted depletion of *pks+ E. coli* from the gut microbiome using specific antibodies^54^ or through innovative engineered phage therapy^55^. This is particularly important for people at risk for developing CRC, such as individuals with a compromised intestinal epithelial or mucus barrier, IBD patients and people with genetic predisposition for CRC development such as familial adenomatous polyposis (FAP) and Lynch-syndrome patients.

In conclusion, using the Zeb2^IEC-Tg/+^ microbiota-dependent mouse CRC model, we have demonstrated for the first time that bacterial FimH/FmlH-dependent adhesion to the host epithelium promotes colibactin-mediated genotoxicity by *pks+ E. coli*. Targeting adhesion of *pks+ E. coli* and other *Enterobacteriaceae* could thus be an effective strategy to restrain the pro-tumorigenic effects of colibactin. Our study also underscores the need for prudence when considering *pks+* probiotic strains, as the uncertain risk posed by gain-of-function mutants could elevate the inherent genotoxic activity of these strains.

## Method details

### Bacterial strains, growth conditions and cloning

Bacteria were grown under type 1 pili inducing conditions, as described previously^56^. Briefly, a single colony was inoculated in 5 mL of Luria Broth (LB) and incubated at 37°C under static conditions for 24 h. Bacteria were then diluted (1:100) into fresh LB and incubated at 37°C under static conditions for 18–24 h. Bacteria were subsequently washed three times with PBS and then concentrated to 1×10^8^ CFU per 100 μL.

Adhesin deletion strains were constructed using the λ red recombination method^57^. The *fimH* or *fmlH* gene was replaced by spectinomycin resistance cassette, using primers with 35 nt homology regions, MK179 and MK180 for *ΔfimH* and MK181 and MK182 for *ΔfmlH*, and a modified pKD46 vector (pKD46_Tmp). *E. coli* Nissle *ΔclbQ* strain was constructed in a similar manner, using primers MK192 and MK193 (Supplementary table 1).

**Supplementary table 1.**
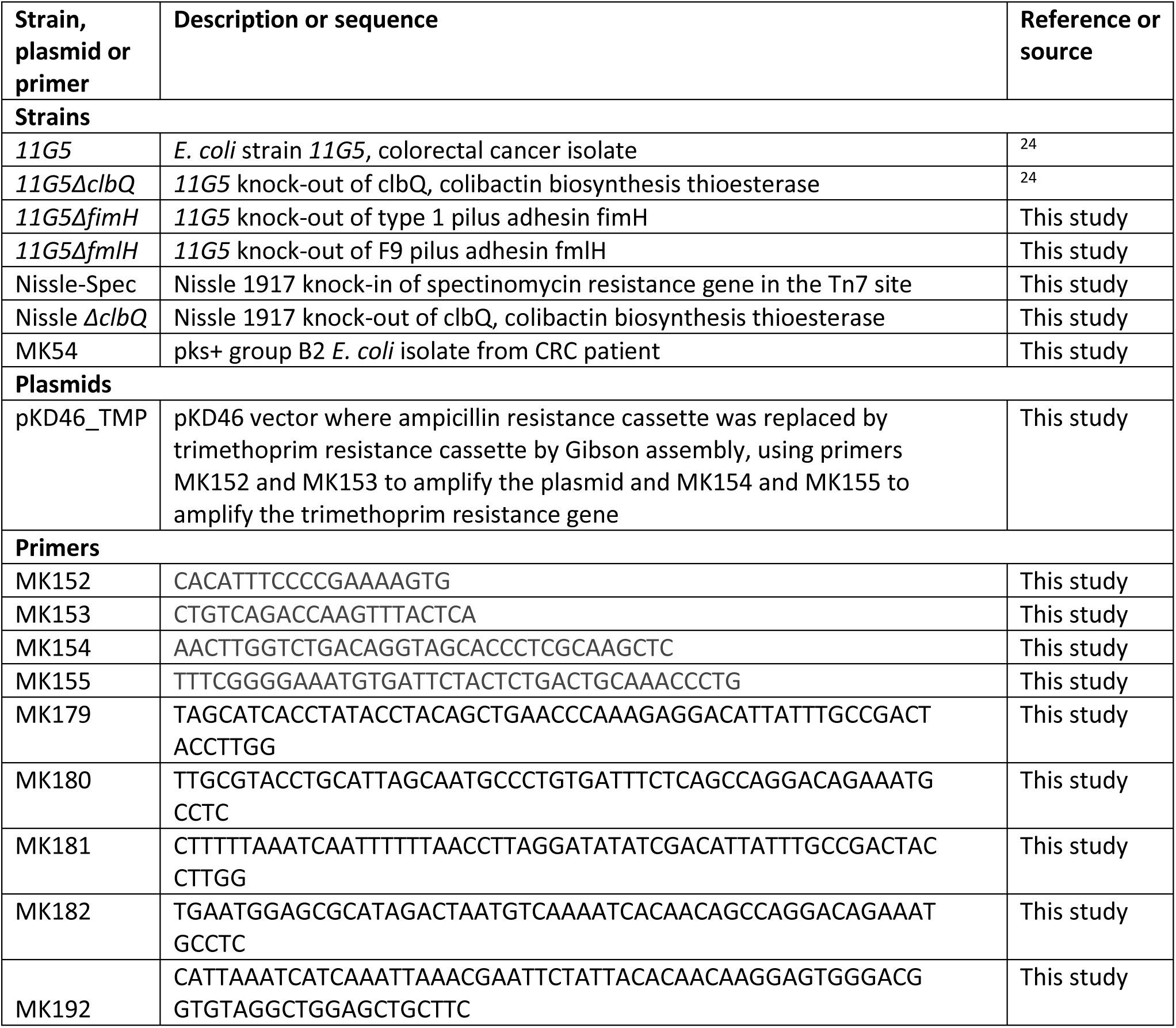
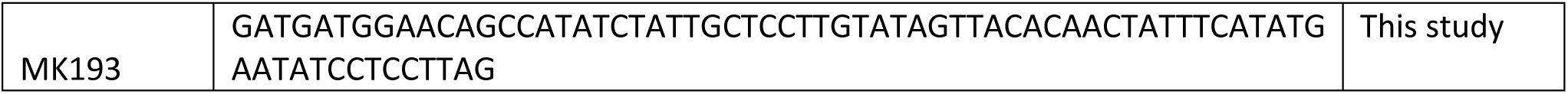
List of strains, plasmids and primers used in this study.

#### Animal experiments

Zeb2^IEC-Tg/+^ mice were described previously. Mice were housed in individually ventilated cages at the VIB Center of Inflammation Research in a specific pathogen-free animal facility. All experiments on mice were conducted according to institutional, national and European animal regulations. Animal protocols were approved by the ethics committee of Ghent University’s Faculty of Sciences.

6-week old Zeb2^IEC-Tg/+^ mice were administered 20 mg streptomycin (Sigma, S1277) by oral gavage 24h before infection. Mice were orally challenged with 10^8^ CFU and followed up for 28 days. Colonisation was followed weekly by plating stool samples.

#### Therapeutic treatment studies

Zeb2^IEC-Tg/+^ mice were infected with 10^8^ CFU *11G5* as described above. 24h after infection, mice were treated daily with 10mg/kg sibofimloc (Medchemexpress, HY-12820) or DMSO vehicle by oral gavage for a period of 4 weeks.

#### Histology

Colon was dissected, fixed in 10% formalin, embedded in paraffin and sectioned at 5 μm. Sections were stained with Hematoxylin/Eosin (HE). For Alcian Blue and Periodic Acid stainings (AB-Pas), dewaxed slides were incubated in 1% Alcian blue for 20 min. Sections were subsequently washed with water and incubated with 1% periodic acid (Sigma, 395132) for 15 min followed by Schiff’s reagent (Sigma, 395-2-016) for 10 min. Sections were counterstained with Mayer’s hematoxylin for 30 s before washing, dehydration and mounting with Entellan (Merck). Sections were imaged on a Zeiss AxioScan Digital Slide Scanner.

#### Histopathological scoring of Zeb2^IEC-Tg/+^ colonic tumor sections

HE and AB-Pas sections were scored blindly according to the scoring system described below (Supplementary table 2). For each tumor, a minimum of 3 sections was scored. For each section, scores were summed resulting in a score ranging from 0 to 16. Scores were averaged per mouse.

**Supplementary table 2.** Histomorphological pathology score of tumors from Zeb2^IEC-Tg+^ mice.

| Score | Immune cell infiltration | Crypt morphology | Mucus aggregates | Invasive lesions into submucosa | Epithelial erosion |
| --- | --- | --- | --- | --- | --- |
| 0 | Normal | Normal | Normal | Normal | Normal |
| 1 | In mucosa | Crypt hyperplasia | Crypt abscesses | ≤ 5 invasive lesions, not transmural | Focal erosion |
| 2 | In mucosa and submucosa | Crypt dysplasia in $\leq 50\%$ of tumor section | $\leq 5$ mucus aggregates | $> 5$ invasive lesions, not transmural | Erosion in $\leq 50\%$ of tumor section |
| 3 | Transmural | Crypt dysplasia in $> 50\%$ of tumor section | $> 5$ mucus aggregates | $>$ transmural invasive lesions | Erosion in $> 50\%$ of tumor section |
| 4 | - | | $> 10$ mucus aggregates | | |

#### Immunofluorescent staining on colon sections

After dewaxing, slides were incubated with antigen unmasking solution (Labconsult, VECH-3300), boiled for 20 min in a PickCell cooking unit and cooled down for 3 h. Sections were blocked in blocking buffer containing 0.25 % goat serum, 0.5 % Fish skin gelatin and 2 % BSA in PBS. For anti-γH2Ax and anti-Epcam staining, sections were blocking in blocking buffer containing 10% goat serum, 0.5 % Fish skin gelatin and 2 % BSA in PBS. Next, slides were incubated with anti-γH2Ax (1:1000, Millipore, clone JBW301), anti-Epcam (1:500, Abcam, ab71916), anti-E-cadherin (1:400, BD, 610182) or *E. coli* (1:250, Abcam, ab 78978) primary antibodies overnight at 4 °C. As secondary antibodies goat-anti-mouse DyLight 488, goat-anti-mouse AlexaFluor 568 (1:500), goat-anti-rabbit DyLight 488 (1:1000) or and goat-anti-rabbit DyLight 633 (1:1000) were used in combination with DAPI.

#### γH2Ax quantification

γH2Ax-Epcam staining was imaged on a Zeiss AxioScan Digital Slide Scanner. For each mouse, gH2Ax positive cells were quantified on three different images. Per image, γH2Ax+ cells were counted in 3 different sections, each section consisting of at least 10 consecutive crypts.

#### Quantification of serum cytokines

Cytokine concentrations in serum were determined by a magnetic bead-based multiplex assay using Luminex technology (BioRad), according to the manufacturer’s instructions.

#### Lipocalin2 ELISA

Fecal lipocalin2 levels were analysed using the mouse Lipocalin2/NGAL ELISA kit (Rndsystems, DY1857) according to the manufacturer’s instructions.

#### Isolation mucosa-associated and crypt-associated bacteria

Following a 4-week infection, mice were sacrificed and colon was dissected and flushed with 1x sterile ice cold PBS. A piece of tumor was opened longitudinally and incubated with 0.016% w/v dithiothreitol solution (30min, 4°C). Supernatant was plated and incubated overnight at 37 °C onto LB agar containing ampicillin (100 μg/ml) for WT *11G5* or spectinomycin (100 μg/ml) for *Nissle 1917*. The next day, mucosa-associated bacterial counts were recorded. To isolate crypt-associated bacteria, crypts were isolated as previously described^58^. Crypts were plated, incubated overnight and enumerated the next day.

#### Isolation of Epcam and CD45 cells for bulk RNA sequencing

WT and Zeb2^IEC-Tg/+^ mice were infected with *E. coli Nissle 1917* or *E. coli 11G5* as described above. Following a 4-week infection, mice were sacrificed, colon was dissected and flushed with 1x PBS. Tumors were dissociated using a mouse tumor dissociation kit (Miltenyi, 130-096-730) according to the manufacturer’s instructions. Total viable cells were sorted on BD FACS Symphony S6 using following antibodies: epithelial cells – CD326 PerCP-eFluor710 (ThermoFisher, catalog #46-5791-82); CD45+ immune cells – CD45-PE (ThermoFisher, catalog #12-0451-81).

#### Transcriptome analysis

RNA was extracted from sorted cells by Trizol (Invitrogen). RNA quality was checked on RNA 6000 nano-chip (Agilent). The library prep, using 12 ng of input RNA, was constructed using the QuantSeq 3’ mRNA-seq FWD kit (Lexogen) and the UMI Second Strand Synthesis Module (Illumina) according to the manufacturer’s instructions. Libraries were amplified with 17 PCR cycles and purified using Purification Module with Magnetic Beads (Lexogen). Final library QC was performed on a Bioanalyzer using a High Sensitivity DNA chip (Agilent) and via RT-qPCR according to the Illumina Sequencing Library qPCR Quantification Guide. The libraries were sequenced as single-end 76 using a NextSeq500 device (Illumina). Sequencing quality of raw FASTQ reads were assessed by FastQC (v0.11.9) and MultiQC software (v1.12)^59,60^. FASTQ reads were subsequently quality-filtered using Trimmomatic (v0.39)^61^. Transcripts were mapped against *Mus musculus* (C57BL/6 mouse) reference genome GRCm39 using STAR (v2.7.10a)^62^. BAM files were created using Samtools (v1.9) and counted with featureCounts from the Subread package (v2.0.3)^63,64^. The experimental design was assessed using principal component analysis (PCA) using DESeq2’s plotPCA function. Graphs were plotted in R using ggplot2 (v3.4.1)^65^. Differential Gene Expression (DGE) analysis was performed using DESeq2 (v1.38.2). P-values were adjusted using Benjamini-Hochberg correction and genes were labeled as differentially expressed when the adjusted-p value < 0.05. Functional enrichment was performed using the fgsea (v1.24.0)^66^. Pathways were obtained from the Molecular Signatures Database (MsigDB)^67,68^. DGE results were shrunk using the ashr package in R (v2.2-54) to account for lowly expressed genes having a large dispersion^69^. A preranked genelist was generated by ordering the shrunk DGE results according to log_2_FC. Enrichment results were filtered using a normalized enrichment score (NES) cut-off > 1. To investigate reactome pathways linked to DNA damage, functional enrichment was performed using genekitr (v1.1.0). Gene set variation analysis (GSVA) (v1.46.0) was used to generate GSVA enrichment scores and were visualized using pheatmap (v1.0.12)^70,71^.

#### Cell culture

HT-29 and HCT116 cells were cultured in complete DMEM (Gibco) supplemented with 10% fetal calf serum (Tico), 1% L-Glutamine (Lonza), 0,4% sodium-pyruvate (Sigma) and 1% non-essential amino acids (Lonza) at 37 °C in a 5% CO_2_ humidified environment.

#### *In vitro* infection

Cells were infected with a ratio of 100 *E. coli* to 1 cell (multiplicity of infection MOI 100), spun at 300xg for 1 min to maximize bacterial:epithelial cell contact, and infected for indicated timepoints. For γH2Ax analysis: after a 6h infection at 37 °C and 5% CO_2_, cells were washed 3 times with 1x PBS and incubated until analysis in cell culture medium supplemented with gentamicin (Sigma, G1272). As positive control, cells were treated with 1 uM staurosporine (Abcam, ab120056).

#### Western blot analysis

After infection, cells were lysed in NP-40 buffer (50 mM Tris-HCl, pH 7.6; 1 mM EDTA; 150 mM NaCl; 1% NP-40; 0.5% sodiumdeoxycholate; 0.1% SDS) containing protease inhibitors (Roche) and phosphatase inhibitors (Sigma), denatured in 1 x Laemmli buffer (50 mM Tris-HCl pH8.2; 2% SDS; 10% glycerol; 0.005% BFB; 5% β-mercapto-ethanol) and boiled for 10 min at 95 °C. 40 ug of cell lysates were separated by by SDS-polyacrylamide gel electrophoresis (PAGE), transferred to nitrocellulose and analysed by immunoblotting. Membranes were probed with antibodies against γH2Ax (1:1000, Sigma, clone JBW301) and actin-HRP (1:10000, MP Biomedicals). As secondary antibody, HRP-coupled anti-mouse-HRP was used (1:2500, Amersham) and detection was done by chemiluminescence (Western Lightning Plus ECL, Perkin Elmer) using the Amersham Imager 600 (GE Healthcare).

#### Immunofluorescence on cells

HT-29 cells were infected at MOI of 100 as described above. After a 3h infection, cells were fixed in 4% paraformaldehyde for 10min at RT. After washes, cells were permeabilized in 0.2% Triton X-100 and blocked in PBS containing 0.5% BSA, 0.02% TX-100 and 1% goat serum. Primary antibodies were incubated overnight at 4 °C in blocking buffer. Secondary antibodies used are listed in Supplementary table 2 in combination with DAPI.

#### Flow cytometry-based adherence assay

Flow cytometric analysis of bacterial adhesion was performed as previously described^72^. Briefly, *E. coli* cultures were stained with CFSE (Cell-Trace CFSE Cell Proliferation Kit, Thermofisher, C34554) before infection of HCT116 and HT-29 cells at MOI 100, spun down for 4 min 300 x g to synchronize adhesion and incubated at 37°C, 5% CO_2_ for 3h. Cells were washed 3x with 1x sterile PBS to remove non-adherent and loosely-adhered bacteria. Trypsin was added to each well, cells were centrifuged for 10 min at 1000 x g and then resuspended in PBS for flow cytometric analysis. All experiments were carried out in triplicate and at least 100.000 events were recorded with a flow cytometer.

#### Quantification bacterial adhesion to epithelial cells *in vitro*

To quantify bacterial adhesion to *in vitro* epithelial cell cultures, *in vitro* adhesion assays were performed as previously described^73^. Briefly, HT-29 cells were infected with *E. coli* strains at MOI 100 for 3h. For determining total number of bacteria (adherent + nonadherent), medium containing bacteria was not taken off the cells, but was lysed directly with 1% Triton X-100. For adherent bacteria, medium was taken off and cells were washed 3x with 1x sterile PBS before lysis with 1% Triton X-100. Serial dilutions were plated on LB agar and incubated overnight at 37°C. Colonies were counted the next day.

#### Scanning electron microscopy (SEM)

HT-29 cells were infected at MOI 100 for 3h infection. For in vivo imaging, colons were dissected and flushed of Zeb2^IEC-Tg/+^ mice infected with *E. coli Nissle* or *E. coli 11G5* for two weeks. Samples were incubated in freshly prepared fixative (2 % paraformaldehyde (EMS), 2.5 % glutaraldehyde (EMS) in 0.1 M Sodium Cacodylate (EMS) buffer, pH7.4) overnight at 4 °C. Fixative was removed by washing 5 x 3 min in 0.1 M cacodylate buffer and samples were incubated in 2% osmium (OsO4, EMS) in 0.1 M cacodylate buffer for 30 min at RT. After washing in H2O for 3 x 5 min, the samples were dehydrated using solutions of increasing EtOH concentration (50 %, 7 0%, 85 %, 9 5%, 2x 100 %), for 15 min each. The samples were then dried in a critical point dryer (Leica EM CPD300) and mounted on an aluminum stub (EMS) using carbon adhesive tape (EMS). Samples were coated with 10 nm of Platinum (Quorum Q150T ES). Colon samples were coated with 15 nm of Platinum. SEM imaging was performed using a Zeiss Crossbeam 540.

### Gentamicin protection assay and identification of new isolates

Isolation of *E. coli* from the biopsy samples was performed as described previously^74^. Briefly, a piece of snap frozen biopsy tumor sample was rotated in 0.016% dithiothreitol (DTT) for 15 min at RT to remove the mucus layer. The tissue was then incubated in gentamicin solution (50 mg/L) for 30 min at 37°C to kill the extracellular bacteria. After 4 washes, the biopsy was placed in 0.01% Triton-X-100 to lyse cells and release the intracellular bacteria. The cell lysate was plated on MacConkey agar and the plates were incubated overnight at 37°C. Lactose fermenting isolates (i.e. pink colonies) were picked up as candidate *E. coli* strains and confirmed by MultiLocus Sequence Typing (MLST) of *adk*, *fumC*, *gyrB*, *icd*, *mdh*, *purA* and *recA* (Achtman scheme^75^), and the presence of *clb* (pks synthesis) genes. Isolate MK54 used in this study, is pks+ group B2 *E. coli* strain.

### Targeted analysis and quantification of *Escherichia coli* genes of interest

The presence of *E. coli* in TCGA sequencing data was determined by analyzing raw sequencing data from The Cancer Microbiome Atlas (TCMA)^36,76^. First, WGS and RNA-Seq from TCGA were screened for microbial content using PathSeq pipeline^77,78^, available as part of the Broad Institute’s Genome Analysis Toolkit (GATK version 4.0.3). We next performed a targeted analysis by mapping annotated *E. coli* genomes to libraries of putative microbial reads generated from this analysis. Annotated genome sequences for *E. coli* strain K12 (GCA_000005845.2) and the pks gene cluster (AM229678.1) were downloaded from GenBank and mapped to these libraries using STAR^62^ without allowing for spliced alignments (--alignIntronMax=1).

### Quantification and statistical analysis

Results are expressed as mean ± SEM, with n the number of mice. Normality tests were performed to check for normal distribution of the data. Statistical significance between uninfected Zeb2^IEC-Tg/+^ and different infections was assessed using a one-way ANOVA followed by Tukey’s multiple comparison test, unless stated otherwise. To quantify γH2Ax in tumors: γH2Ax+ cells were counted in at least 10 consecutive crypts in three areas of a tumor section. For each mouse, at least 3 sections were quantified. For γH2Ax+ quantifications of 11G5 vehicle and sibofimloc treatments, unpaired t-test with Welch’s correction was performed. Bacterial adhesion on tumor sections was performed in ZEISS Arivis software. Three sections were imaged and quantified per mouse. Analyses were performed with Prism 9 software. Statistical details can be found in the figure legends. Fecal colonisation data was analysed as repeated measurements using method of residual maximum likelihood (REML), as implemented in Genstat version 22. Briefly, a linear mixed model (random terms underlined) of the form log(count) = µ + litter + treatment + time + treatment.time + litter.genotype + litter.genotype.time + subject.time was fitted to the 11G5 vs adhesin mutant strains data; a similar linear model without the litter fixed effect and its random interaction terms was fitted to the 11G5 + sibofimloc data. The random term subject.time represents the residual error term with dependent errors because the repeated measurements are taken in the same individual, causing possible correlations among observations. The autoregressive correlation model was selected as best fitted model based on the Akaike’s information criterion coefficient. The significance of the fixed terms in the model and the changes in differences between treatment across time, were assessed using an approximate F-test as implemented in Genstat version 22.

**Supplementary table 3.**
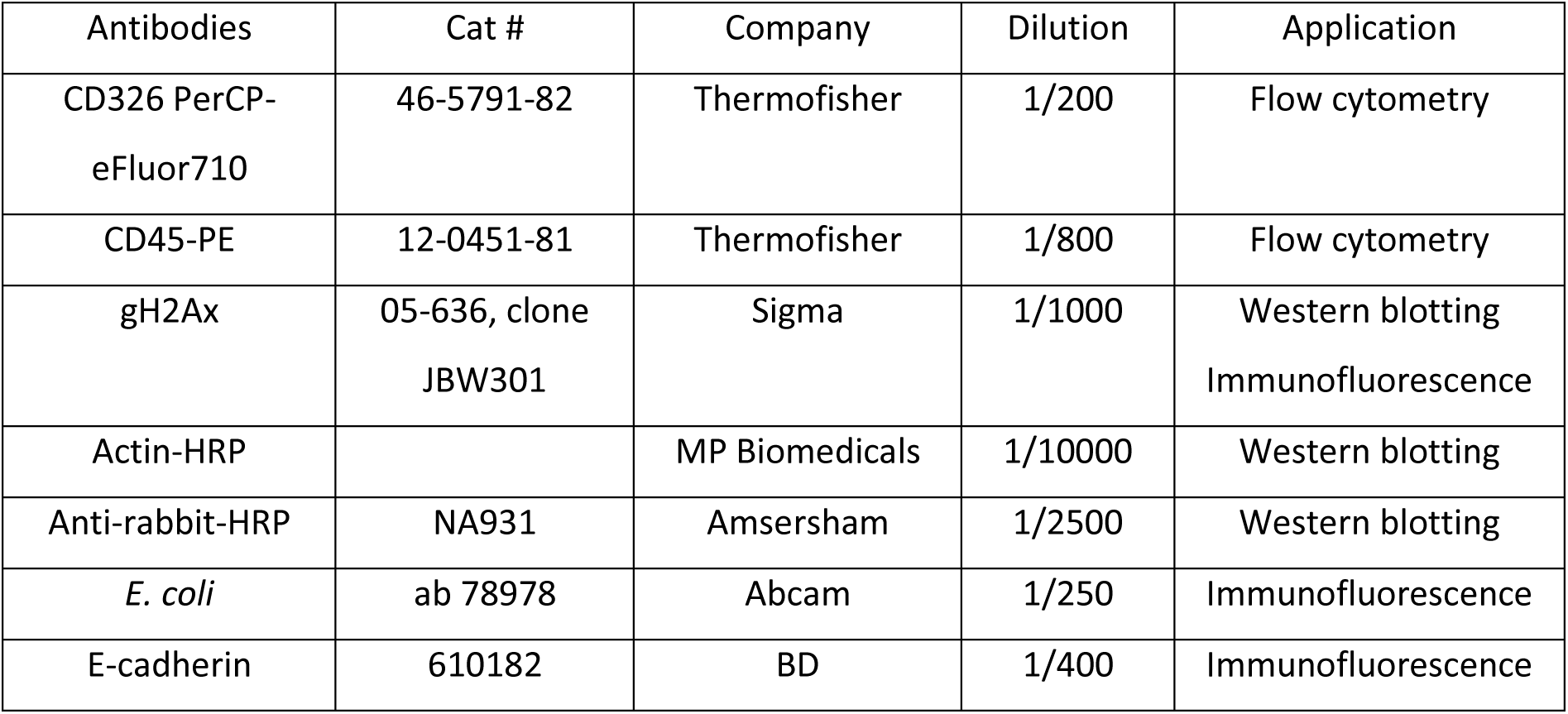
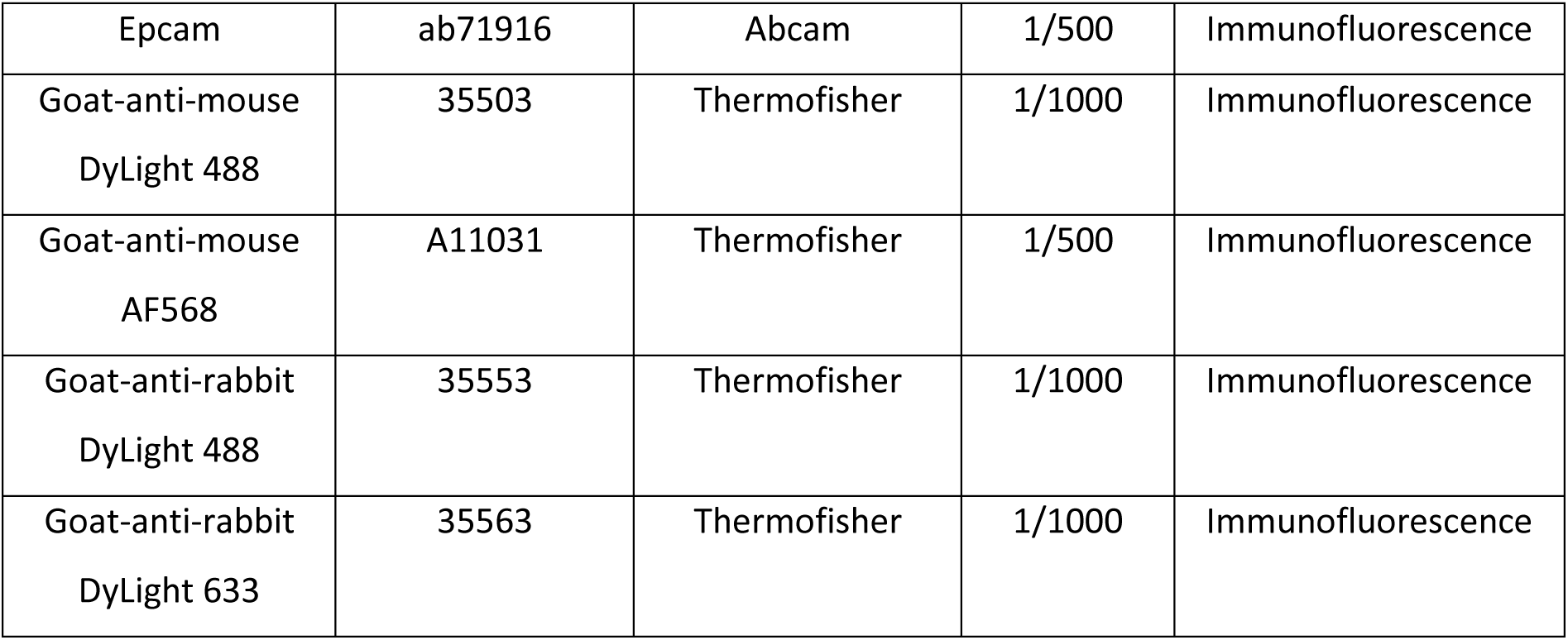
List of antibodies used.

| Antibodies | Cat # | Company | Dilution | Application |
| --- | --- | --- | --- | --- |
| CD326 PerCP-eFluor710 | 46-5791-82 | Thermofisher | 1/200 | Flow cytometry |
| CD45-PE | 12-0451-81 | Thermofisher | 1/800 | Flow cytometry |
| γH2Ax | 05-636, clone<br>JBW301 | Sigma | 1/1000 | Western blotting<br>Immunofluorescence |
| Actin-HRP |  | MP Biomedicals | 1/10000 | Western blotting |
| Anti-rabbit-HRP | NA931 | Amsersham | 1/2500 | Western blotting |
| *E. coli* | ab 78978 | Abcam | 1/250 | Immunofluorescence |
| E-cadherin | 610182 | BD | 1/400 | Immunofluorescence |
| Epcam | ab71916 | Abcam | 1/500 | Immunofluorescence |
| Goat-anti-mouse<br>DyLight 488 | 35503 | Thermofisher | 1/1000 | Immunofluorescence |
| Goat-anti-mouse<br>AF568 | A11031 | Thermofisher | 1/500 | Immunofluorescence |
| Goat-anti-rabbit<br>DyLight 488 | 35553 | Thermofisher | 1/1000 | Immunofluorescence |
| Goat-anti-rabbit<br>DyLight 633 | 35563 | Thermofisher | 1/1000 | Immunofluorescence |

## AUTHOR CONTRIBUTIONS

M.J., M.K., G. Blancke, M.S. and A.T performed the experiments. M.J., M.K., M.C., G. Berx, A.B.D., T.K., G.v.L., H.R. and L.V. performed data analysis. S.T. provided biopsy samples. G.v.L., I.D.I, H.R. and L.V. provided ideas and coordinated the project. M.J. and L.V. wrote the manuscript.

## ACKNOWLEDGEMENTS

We thank G. Dalmasso and R. Bonnet for the *11G5* and *clbQ* mutant strains. We are grateful to L. Bellen and D. Huyghebaert for animal care. We also thank E. Parthoens, P. Borghraef and F. Baeke from the VIB bioimaging core. Schematic representations were made with Biorender.com. M. Jans is a predoctoral fellow supported by an FWO doctoral fellowship. Research in the L. Vereecke lab was supported by grants from Ghent University (BOF.GOA031-22, BOF.IBF037-20), the FWO (EOS-G0H2522N-40007505) and Foundation against Cancer (F/2020/1421). Research in the G. van Loo lab was supported by VIB and research grants from Ghent University (BOF/24J/2021/052 and BOF23/GOA/001) the FWO (G090322N, G026520N, G012618N, EOS-G0H2522N-40007505), the Queen Elisabeth Medical Foundation, the Charcot Foundation, the “Belgian Foundation against Cancer” (F/2018/1200 and F/2022/1899), and the FOREUM Foundation for Research in Rheumatology.

I.D.I. was supported by R01DK113136, R01DK121977, R01AI163007 and the Cancer Research Institute Lloyd J. Old STAR Award.

