## Supplementary figures for "Colibactin-induced genotoxicity and colorectal cancer exacerbation critically depends on adhesin-mediated epithelial binding"

### Supplementary figure legends

**SUPPLEMENTARY FIGURE 1. *E. coli 11G5* infection does not increase colonic inflammation in Zeb2$^{IEC-Tg/+}$ mice and does not affect epithelium in WT littermates. A.** Fecal lipocalin2 levels of uninfected (n = 6), *Nissle 1917*- (n = 5) and *11G5*-infected (n = 5) Zeb2$^{IEC-Tg/+}$ mice following a 4 week infection. Data are represented as mean ± SEM (Kruskal-Wallis with multiple comparisons). **B.** Levels of IL-17A, CXCL-1, IL-1b,IL-6, TNF-a and IFNg in serum of Zeb2$^{IEC-Tg/+}$ mice following a 4 week infection with *Nissle 1917* (n = 4) or *11G5* (n = 8) or uninfected control (n = 3). Data are represented as mean ± SEM (* p < 0.05, *** p<0.001, **** p<0.0001; one-way ANOVA with multiple comparisons). **C.** Principal component analysis (PCA) showing no changes in overall gene expression profiles in CD45+ immune cells sorted from the colon of Zeb2$^{IEC-Tg/+}$ mice following 4-week *Nissle 1917* (n = 5) or *11G5* (n = 4) infection. **D.** Volcano plot of differentially expressed genes (DEG) in colonic CD45+ immune cells of *11G5* infected Zeb2$^{IEC-Tg/+}$ mice compared to *Nissle 1917* (8 genes downregulated and 2 genes upregulated in *11G5* conditions compared to *Nissle 1917*). **E.** Percentage of bacteria closely associated to the colonic epithelium (minimal average distance to epithelium < 20 µm) and not associated with the epithelium (minimal average distance to epithelium ≥ 20 µm) after *Nissle 1917* and *11G5* infection or uninfected controls. Data are represented as mean ± SEM. (*** p<0.001, **** p<0.0001, two-way ANOVA). **F.** Epithelial surface-associated and crypt-associated *Nissle 1917* and *11G5* in proximal and distal colonic segments of Zeb2$^{IEC-Tg/+}$ mice after a 4 week infection. Data are represented as mean ± SEM (* p < 0.05, ** p<0.01; two-way ANOVA with multiple comparisons). **G**. Heatmap of gene set variation analysis (GSVA) enrichment scores for indicated EMT-associated pathways in colonic Epcam+ cells from *Nissle 1917*- and *11G5*-infected Zeb2$^{IEC-Tg/+}$ mice. **H.** Representative images of H&E-stained colon sections of wild-type (WT) mice following a 4 week infection *11G5*. Scale bar: 200 µm. **I.** PCA showing no changes in overall gene expression profiles in colonic epithelial cells sorted from the colon of WT mice following 4-week *Nissle 1917* (n = 3) or *11G5* (n = 4) infection. **J.** Volcano plot of DEG in colonic epithelial cells of *11G5* infected WT mice compared to *Nissle 1917* (8 genes downregulated and 26 genes upregulated in *11G5* conditions compared to *Nissle 1917*). **K.** Volcano plot of DEG in colonic CD45+ cells of *11G5* infected WT mice compared to *Nissle 1917* (4 genes downregulated in *11G5* conditions compared to *Nissle 1917*).

**SUPPLEMENTARY FIGURE 2 – *E. coli 11G5ΔclbQ* colonizes similarly as WT *11G5* and is adhesion-competent. A.** Fecal colonisation at sacrifice. Data are represented as mean ± SEM (unpaired t-test). **B.** Representative images of *E. coli* (green) and Epcam (red) immunostaining of tumor sections of *11G5* or isogenic colibactin mutant (*11G5ΔclbQ*) infected Zeb2$^{IEC-Tg/+}$ mice following a 4-week infection (lower panel: close-up). Scalebar overview, 50 µm; scalebar zoom, 20 µm.

**SUPPLEMENTARY FIGURE 3 – Isogenic adhesin mutant strains *11G5ΔFimH* and *11G5ΔFmlH* are capable of colonizing Zeb2$^{IEC-Tg/+}$ mice but no longer adhere to and induce DNA damage response in HCT-116 and HT-29 cells *in vitro*. A.** Fecal colonisation of WT 11G5 and adhesin mutant strains monitored over time. Data are represented as mean ± SEM. **B & C.** Cecal and colonic colonisation after 4-week infection. Data are represented as mean ± SEM. (* p < 0.05; one-way ANOVA with multiple comparisons) **D.** Representative scanning electron microscopy (SEM) images of HT-29 cells infected with E. coli 11G5 and clbQ isogenic mutant. Infection of 3h. Scale bar: 5 µm. **E.** Schematic representation of flow cytometric analysis of bacterial adhesion. Bacteria are CFSE-labeled and added to epithelial cell culture. **F.** CFSE-gate on HCT-116 cells infected for 3h with CFSE-stained *E. coli* strains. **G.** Flow cytometric analysis of bacterial adhesion on HT-29 cells after 3h infection. Data are represented as mean ± SD and is representative of three independent experiments (**** p<0.0001; one-way ANOVA with multiple comparisons). **H.** Western blot analysis of γH2Ax on HCT116 cells infected with *Nissle 1917*, *11G5* or isogenic mutants. Actin was used as loading control. Data is representative for three independent experiments. **I.** Immunofluorescent staining for γH2Ax in HT-29 cells infected with *E. coli Nissle 1917*, *11G5* or isogenic mutants. Cells were infected for 6h and medium was replaced with gentamicin-containing medium for 18h. Scale bar: 10 µm.

**SUPPLEMENTARY FIGURE 4 – new *E. coli* strain isolated from tumor biopsy accelerates tumor development *in vivo* while sibofimloc reduces epithelial binding of and abrogates their DNA damaging capacity *in vitro*. A.** Flushed colon weights of Zeb2$^{IEC-Tg/+}$ mice after *Nissle 1917* (n = 4) and *pks+ E. coli strain MK54* (n = 4). Data are represented as mean ± SEM. (* p < 0.05Mann-Whitney test). **B.** Representative images of hematoxylin eosin (H&E) stained colon sections of Zeb2$^{IEC-Tg/+}$ mice after *E. coli Nissle 1917* and *pks+ E. coli MK54* infection. Scale bar: 200 µm. **C.** Histopathological score of tumor sections following *MK54* and *Nissle 1917* infection. Data are represented as mean ± SEM. (**p < 0.01; two-sided unpaired t-test). **D.** Mean number of invasive lesions into the submucosa. Data are represented as mean ± SEM. (*p < 0.05; two-sided unpaired t-test). **E.** Representative images of *E. coli* (green) and E-cadherin (red) staining on tumor sections of *Nissle 1917*- or *MK54*-infected Zeb2$^{IEC-Tg/+}$ mice following a 4-week infection. Scale bar 50 µm. **F.** Percentage of bacteria closely associated to the colonic epithelium (minimal average distance to epithelium < 20 µm) and not associated with the epithelium (minimal average distance to epithelium ≥ 20 µm) after *Nissle 1917* and *pks+ E. coli MK54* infection. Data are represented as mean ± SEM. (two-way ANOVA). **G.** Representative images of γH2Ax (green) and Epcam (red). Scale bar: 100 µm. **H.** Quantification of the number of γH2Ax-positive epithelial cells in tumor sections of Zeb2$^{IEC-Tg/+}$ mice after infection. Data are represented as mean ± SEM (* p < 0.05; two-sided unpaired t-test with Welch's correction). **I.** flow cytometric analysis of bacterial adhesion to HCT116 cells after 1h infection and sibofimloc treatment. Data are represented as mean ± SD and is representative of two independent experiments (**** $p<0.0001$; one-way ANOVA with multiple comparisons). **J.** HCT116 cells were infected with pks+ *E. coli* MK54 (MOI 10) for 1h. Immunoblot analysis for γH2Ax 24h after infection. Actin was used as loading control.

### SUPPLEMENTARY FIGURE 1

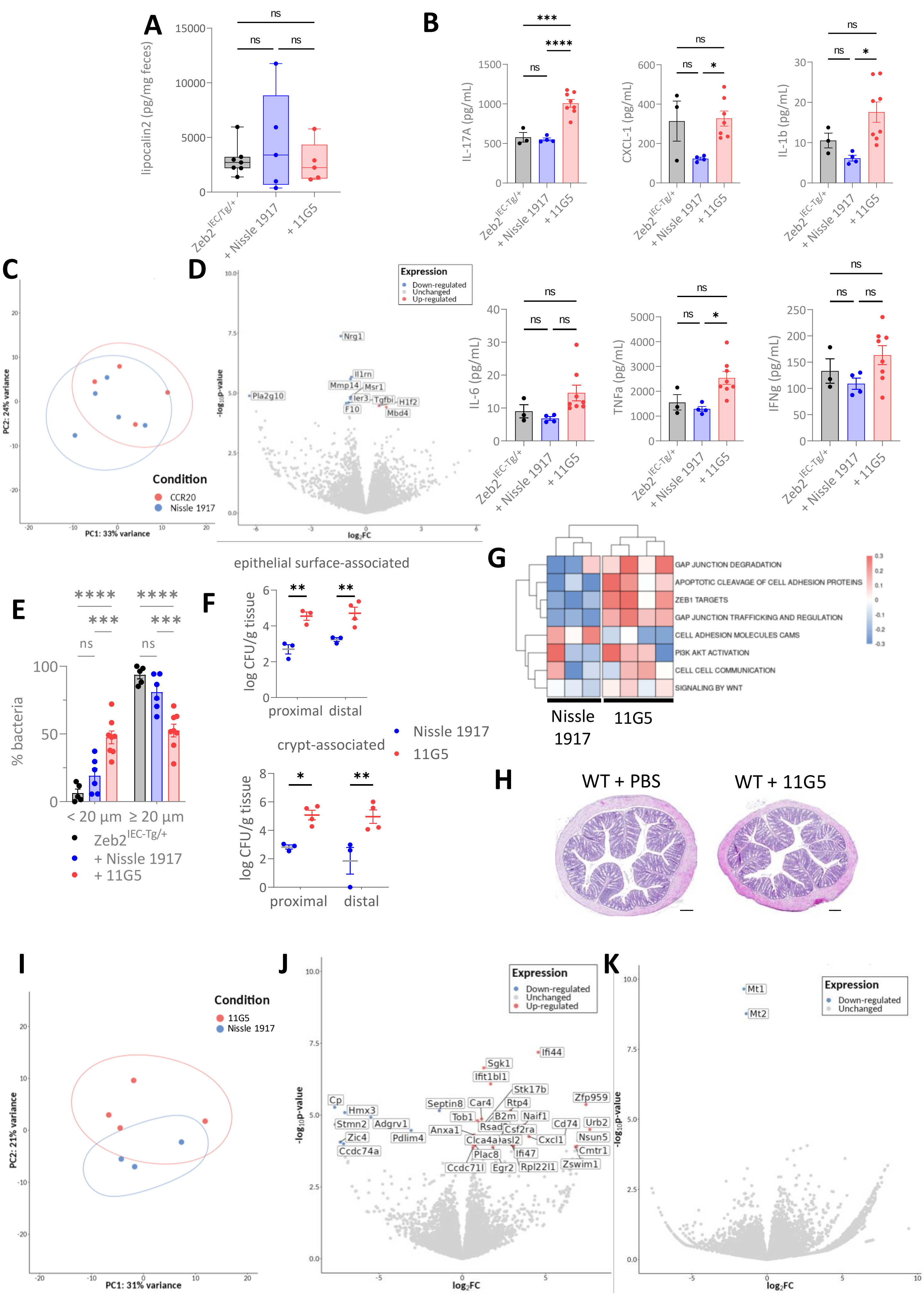

### SUPPLEMENTARY FIGURE 2

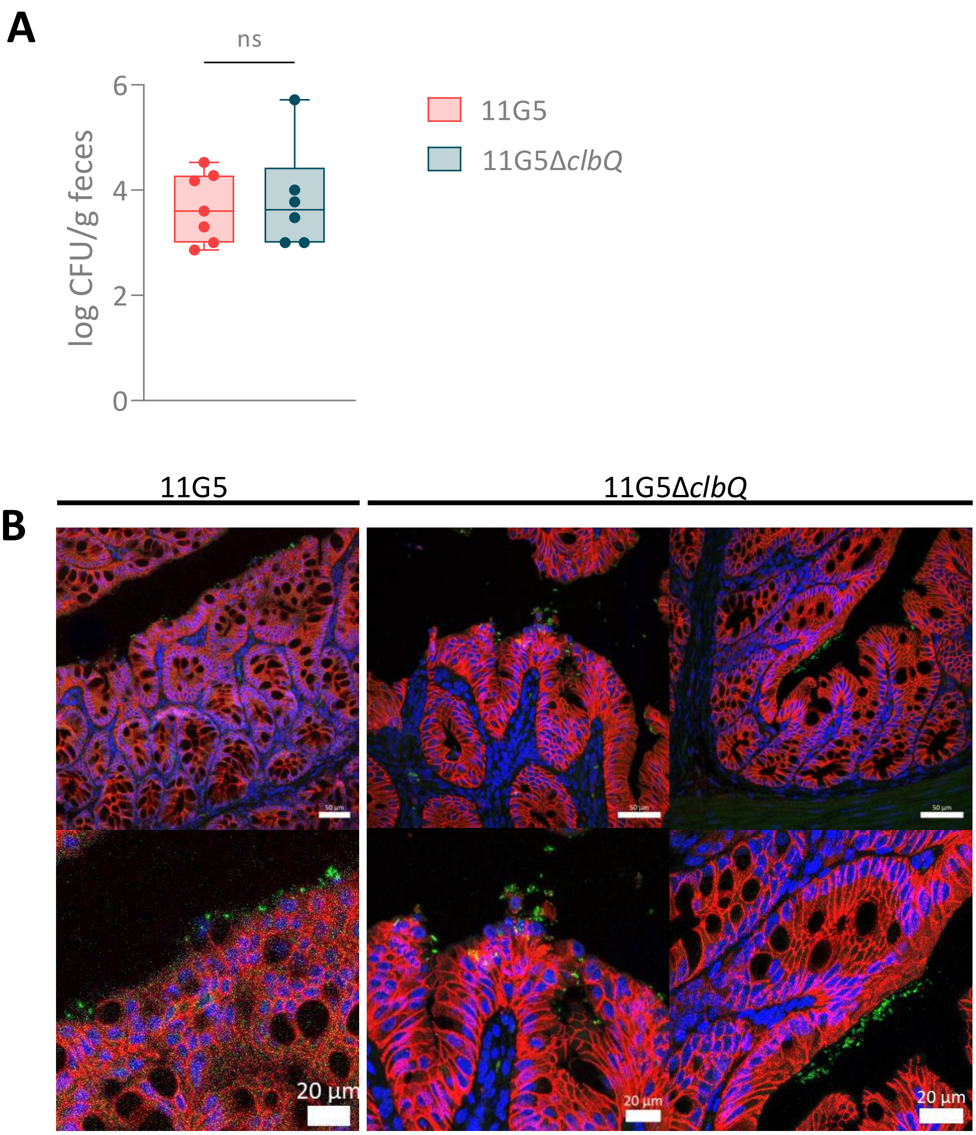

SUPPLEMENTARY FIGURE 3

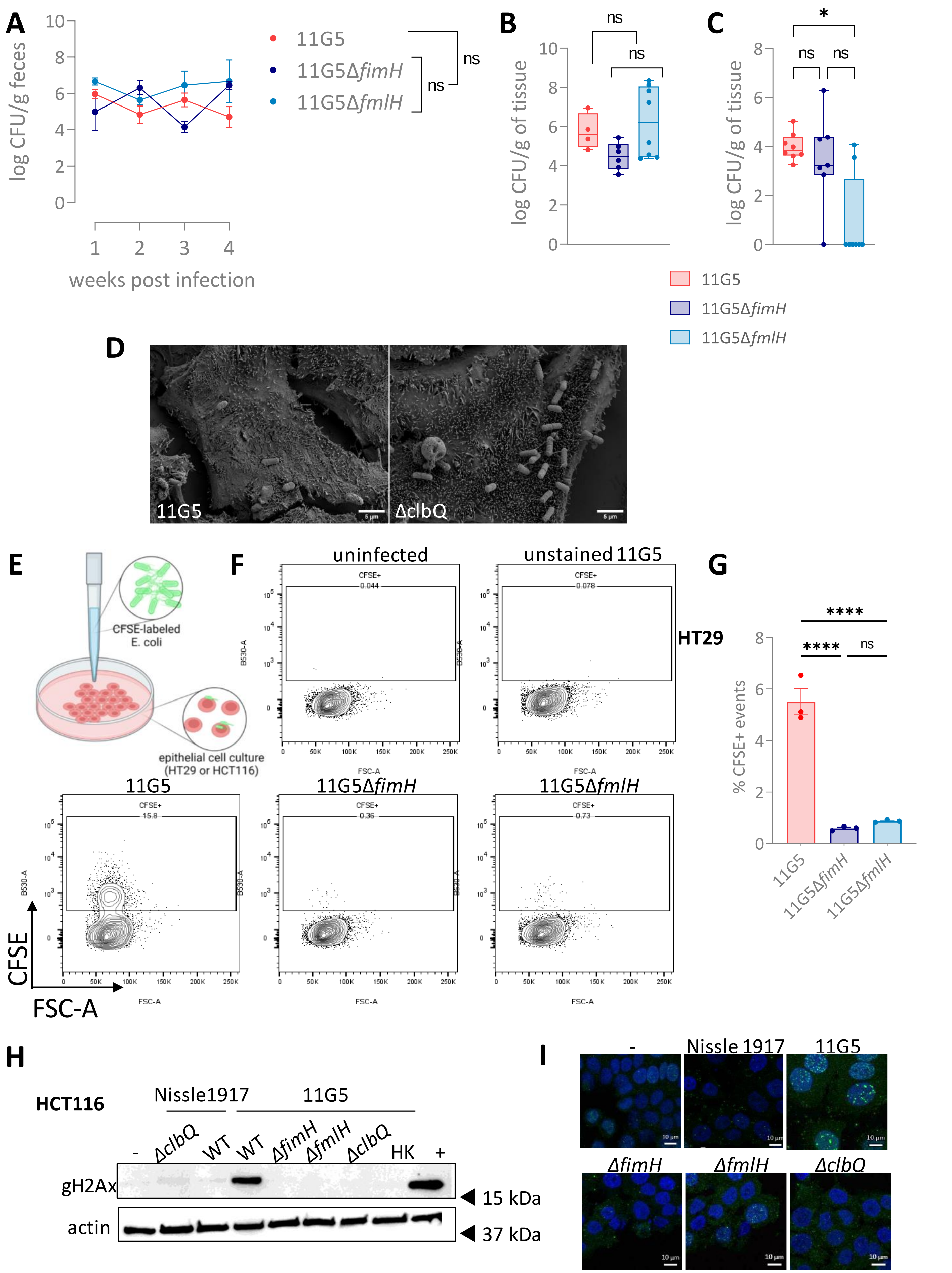

SUPPLEMENTARY FIGURE 4

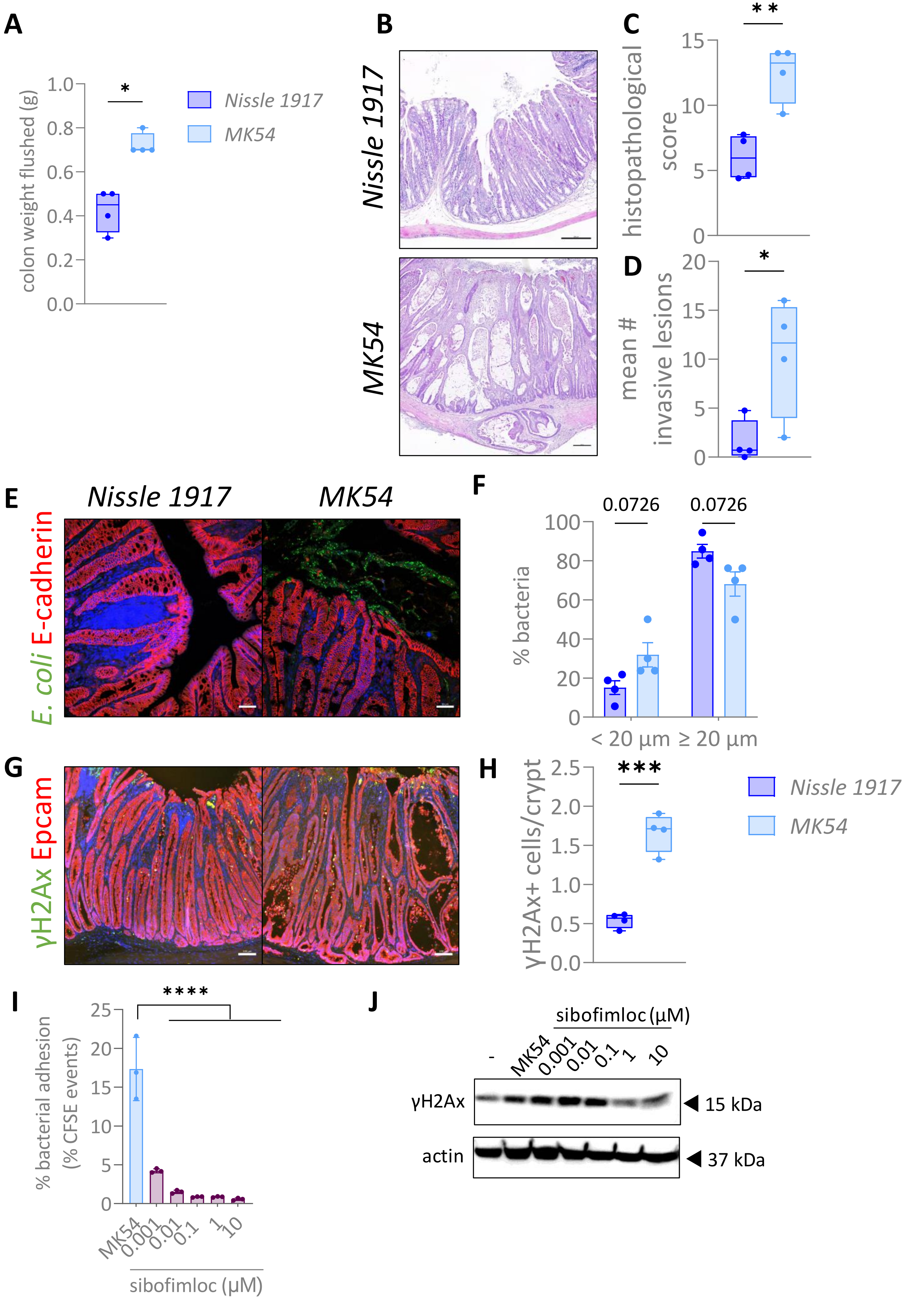
